# Residue-Specific Modulation of Aggregation-Associated Interactions by Spermine in Tau, α-Synuclein, and Aβ40

**DOI:** 10.64898/2025.12.23.696224

**Authors:** Debasis Saha, Sun Xun, Jinghui Luo, Wenwei Zheng

**Affiliations:** College of Integrative Sciences and Arts, Arizona State University, Mesa, AZ, 85212, USA; Center for Life Sciences, Paul Scherrer Institute, Forschungsstrasse 111, 5232 Villigen PSI, Switzerland; Center for Biological Physics, Arizona State University, Tempe, AZ, 85281, USA

## Abstract

Preventing neurodegenerative diseases associated with intrinsically disordered proteins (IDPs) remains a major challenge due to the lack of a detailed, sequence-level picture of disease-relevant structures formation and the influence of cellular factors that modulate these transitions. Here, we probe spermine (Spm), a +4 charged polyamine abundant in cells, to determine how it reshapes the conformational ensembles and fibril-associated contact propensities of three disease-linked IDPs: the K18 domain of Tau, α-synuclein (αS) and amyloid-β40 (Aβ40). Using long all-atom molecular dynamics simulations across a range of Spm concentrations, we quantify residue-level changes in intra-chain contacts relative to native contacts observed in fibrils, and corroborate computational predictions with ThT fluorescence assays for Tau constructs. Spm acts in a sequence-and region-specific manner, not simply through overall net charge. In K18, Spm binds near the fourth microtubule binding repeat, disrupting intra-chain contacts associated with Alzheimer’s fibril structures and thereby inhibiting aggregation. In αS, Spm binds mainly to acidic residues in the C-terminal half of the sequence and redistributes intramolecular contacts in a way that increases contact propensity in the central aggregation-prone region and therefore aggregation, in line with previous studies showing Spm-enhanced αS aggregation. For Aβ40, Spm neutralizes acidic residues near positions 22–24 and shifts the balance of intra-chain interactions toward its aggregation-prone core, resulting in a net promotion of fibril-like conformations. These divergent effects show that net charge alone cannot predict polyamine influence on IDPs. Instead, residue-specific binding hotspots and local reweighting of aggregation-linked contacts determine whether Spm promotes or suppresses fibril-like conformations. This combined simulation–experimental framework provides a mechanistic basis for how small molecules reprogram IDP conformational ensembles and suggests principles for designing ligands that exploit similar residue-level modulation.

## Introduction

The rising prevalence of age-related neurodegenerative diseases among the elderly represents a persistent socio-economic challenge. These conditions are often marked by the intracellular or extracellular aggregation of various disordered proteins, such as Tau or amyloid-β (Aβ), in the case of Alzheimer’s disease (AD),^1^ and α-Synuclein (αS) in Parkinson’s disease (PD). Tau, a microtubule binding protein consisting of 441 residues, exhibits six isoforms in the human brain, with three isoforms containing three microtubule binding repeats (3R) and the other three containing four repeats (4R)^1^. The presence of different isoforms in Tau filaments can lead to distinct Tau folds, contributing not only to AD but also to conditions like Pick’s disease^2^ or familial British dementia^3^, among others. Similarly, different assembly modes of the 140-residue αS leads to diseases such as multiple system atrophy (MSA)^4^ or dementia with Lewy bodies (DLB)^5^. Aβ, another crucial protein implicated in AD^6^, comprises 39-43 residues and typically adopts a random coil structure in solution^7^. However, in AD patients, Aβ peptides form fibrils characterized by stacked β-sheets^8^. The terms “tauopathies” and “synucleinopathies”^9^ have been introduced to categorize diseases resulting from abnormalities in Tau or αS, which can stem from various factors including posttranslational modifications^10^ or specific mutations^11,12^, or through changes in environment.

The disease related fibril formation for Tau, Aβ and αS often takes place via the formation of structurally diverse oligomer formation^13–15^. The transient nature of these oligomers make their characterization challenging, which in turn complicates the identification of suitable molecules capable of preventing protein aggregation. Additionally, substantial structural polymorphism even in their aggregated states^16^ make them largely inaccessible to conventional structural biology methods,^17,18^ leading them being frequently labelled “undruggable”.^19^ Efforts have been made to focus on specific segments on the fibril structure. For example, characterization of fibril structure formed by the residues VQIVYK from the third repeat (R3)^20^ of Tau has enabled the development of inhibitors that prevent Tau aggregation.^21,22^ However, the inhibitors developed based on the particular residue segment has been found to be incapable of preventing aggregation of full length Tau.^22^ Although different studies have been carried out to develop ligands capable of preventing Tau aggregation^23,24^, the concentration required for their effectiveness remains high, making them clinically less effective. Consequently, alternative strategies that shift the conformational landscape of these proteins away from aggregation-prone states represent a promising avenue for therapeutic intervention in neurodegenerative disorders.

In this context, spermine (Spm) represent an intriguing modulator. Among the different naturally occurring polyamines involved in crucial biological processes within cells^25,26^, Spm possesses the highest charge of +4 and is known to be most effective for condensation of other biomolecules.^27^ Spm and spermidine has been found to contribute to neuroprotection by stimulating autophagic pathways that degrade damaged organelles and toxic protein aggregates.^28^ The role of spermine and other polyamines in the aggregation behaviour of Tau, αS and Aβ has been well-studied over the years.^29–31^ While the presence of Spm has been shown to decrease lag and transition times of the aggregation process for both αS^30^ and Aβ^31^, higher-order polyamines has been shown to prevent fibrillization of Tau.^32^ These opposing effects have motivated therapeutic interest in modulating polyamine metabolism, particularly as a means of mitigating Tau aggregation.^33^ A molecular-and sequence-level understanding of these divergent outcomes is therefore essential to elucidate how Spm reshapes the conformational ensembles and aggregation-prone features of these disease-associated proteins.

In this study, we address the Spm mediated conformational modification of the microtubule binding domain of Tau (K18), αS and the 40-residue variant of Aβ (Aβ40) by combining all-atom molecular dynamics (MD) simulations with fluorescence-based validation experiments. Due to their inherent structural heterogeneity, simulating these IDPs at atomic resolution is challenging—particularly in the absence of enhanced sampling techniques^34,35^. Nevertheless, we conduct long-timescale simulations to capture the range of conformations accessible to these proteins. Evidences have shown that intra-chain hairpin formation propensity and aggregation kinetics are correlated for Tau,^36^ underscoring the critical role of intra-chain contacts in driving disease-relevant filament formation. Hence, our approach focuses on whether Spm reshapes intramolecular contacts that correlate with aggregation propensity.

Using fibrillar structures from patient-derived samples as references, we quantify the formation of aggregation-relevant contacts in the presence and absence of Spm. The results reveal that Spm’s effects are not explained by global electrostatics alone; instead, residue-specific interactions dictate whether Spm disrupts or promotes aggregation. We further contextualize our results in terms of liquid–liquid phase separation (LLPS) of these proteins^37^, offering insights into how Spm balance shapes both functional and pathological assemblies. By revealing how Spm reprograms the conformational preferences of IDPs, our study suggests new opportunities for designing ligands that exploit Spm-inspired mechanisms to modulate aggregation.

## Results and Discussion

### Protocol for Investigating Spermine-Mediated Modulation of Disease-Relevant Structures

In this study, we investigate the effect of Spm (structure shown in Fig. S1) on the three disease-relevant IDPs: K18 region of Tau (hereafter K18), αS and Aβ40 using all-atom MD simulations (see Methods). For each system, the details of protein-Spm ratios and their corresponding simulation details such as box size, number of atoms and simulation lengths are given in Table S1 in SI. Additional simulation details are provided in the Methods section. To validate our simulations, we compared calculated ^15^N NMR chemical shifts for K18 (with and without Spm) with experimental data^37^. The chemical shifts were computed using the SPARTA+^38^, based on snapshots extracted from the trajectories. The computed values of ^15^N chemical shifts in both the presence and absence of Spm are shown in Fig. S2 in SI. The overall patterns suggest good qualitative agreement between simulation and experiment, though a few residues exhibited slightly higher deviations between the two conditions, likely arising from the intrinsic uncertainties in calculating chemical shifts from protein conformations.

Following the simulations, we have investigated Spm’s ability to promote or demote several disease relevant structure formation that are associated with these proteins. The protocol used for this purpose has been schematically demonstrated in Fig. 1. At first, the proteins and Spm has been combined at different ratios and the ensemble has been obtained using the MD simulations, as can be seen from Fig. 1A. From these ensembles, the differences in intra-residue contact map has been obtained from simulations without Spm to check which contacts increase or decrease with Spm (Fig. 1C). Simultaneously, different fibril structures associated with these proteins have been collected to obtain the native contacts within the disease relevant structures (Fig. 1B). The presence of diverse filament morphologies in Tau^39^, αS and similar proteins indicates that multiple aggregate conformations can exist. Moreover, both Tau and αS filaments from patients has been found to promote disease relevant fibril formation more effectively than in vitro–assembled filaments^40^, underscoring the biological relevance of disease-specific polymorphs. To capture this diversity, we selected fibril structures directly derived from patient brain tissues for Tau, αS, and Aβ40. Given the structural heterogeneity observed even among filaments associated with the same disease, we first clustered the collected fibril structures based on shared intra-chain contacts. A few differently structured Tau filaments are shown in Fig. 1B, which has been used for clustering and finding the native contacts along with several other reported structures. One such contact map corresponding to a Tau fibril structure is shown in Fig. 1D.

**Figure 1.**
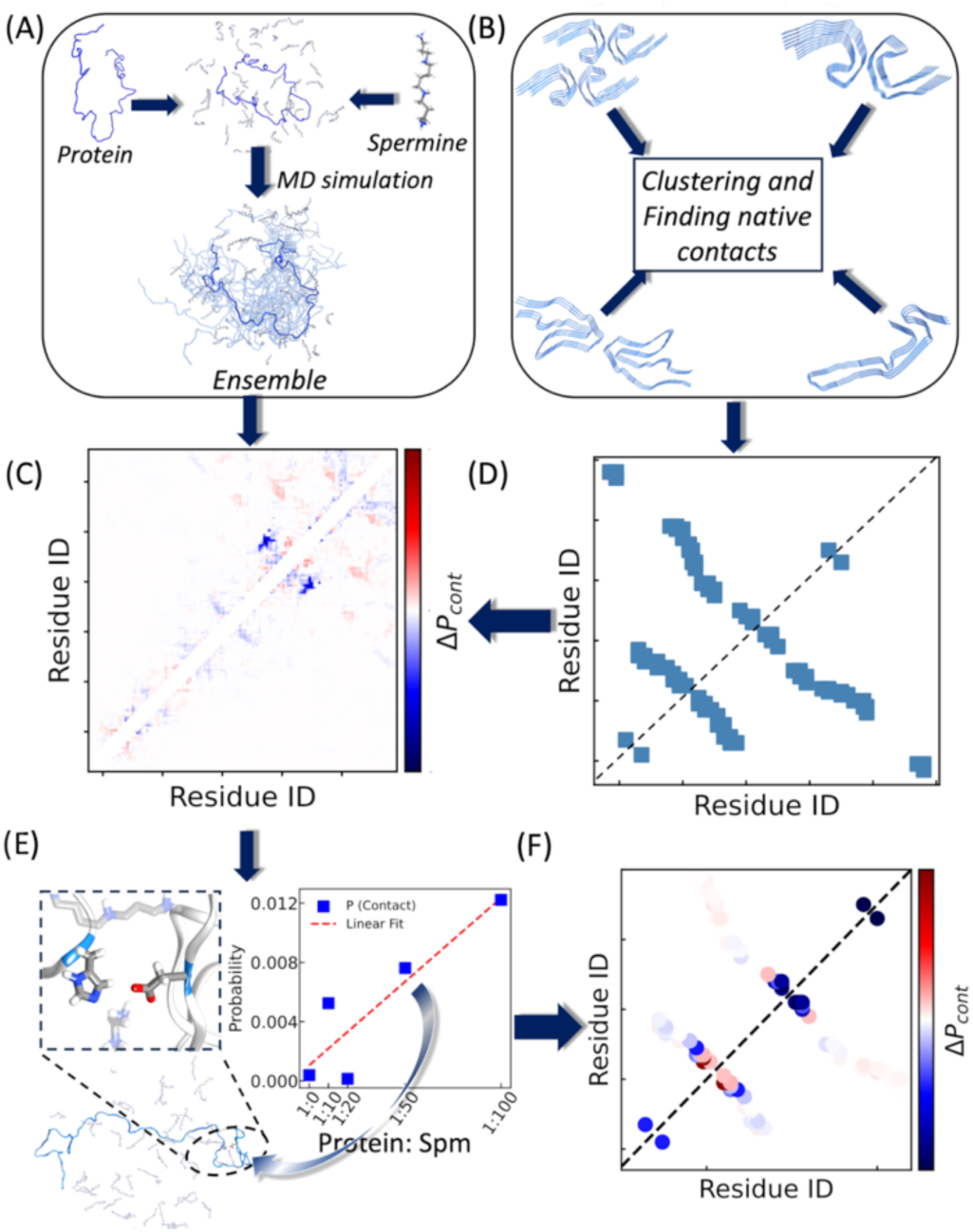
Schematic diagram of the protocol to investigate influence of spermine (Spm) on disease-relevant fibril structures. In the panel (A), IDPs are combined with Spm, and molecular dynamics simulations are performed to generate the structural ensemble. From these ensembles, changes in protein contact maps between the absence and presence of Spm are calculated (panel C). The panel (B) shows representative Tau fibril structures, which are clustered to identify native contacts. One example of contact map from a fibril structure is shown in panel (D), highlighting residue pairs involved in fibril formation. In panel (E), one such contact between residues 330 and 359 observed in the simulation of K18 is illustrated, along with its probability at different Spm ratios. Using this approach, the probability of every fibril contact pair is quantified, producing the distribution shown in panel (F).

For K18 of Tau, among the different structures observed at different conditions, we focus on filaments associated with AD^41^, corticobasal degeneration^42^ and Pick’s disease^2^. The Tau inclusions observed in the Cryo-EM structure reported from AD patients^41^, exhibit polymorphism in the paired helical filaments (PHFs) and straight filaments (SF)^43^ whose cores are made up of double helical stacks of C-shaped subunits as seen previously for other tau filaments.^44^ With only a few residues present in these units, large segments form the N-terminal domain (NTD) and C-terminal domain (CTD) of tau remain disordered in these structures and form a fuzzy coat.^45^ However, while for the structure reported for Pick’s disease^2^ does not have the second repeat (R2) repeat and does not form C-shaped subunits, the filament related to CBD^42^ are not found as double helical stacks. The fact that the filaments associated with AD differ in symmetry between PHFs and SFs^41^ further adds to the complexity involved in structures of tau inclusions. Therefore, in addition to the above mentioned structures, other similar structures reported elsewhere^46,47^ are also included for Tau to find the pairs that are present in fibril structures. For αS, the filament structures chosen here are the ones reported from patients having PD^48^ and multiple system atrophy (MSA)^49^. The αS filaments found in these structures and others obtained in vitro^50^ extends from residues 30 to 100. The PDB id’s belonging to Tau and αS used here have been listed in Table S2 and S3 in SI. For Aβ40 too, structural polymorphism becomes evident from the difference in the β-sheet twist found in the filaments obtained in vitro^51^ and in the one found in AD patients^52^. Therefore, we select two structures found from AD patients^52,53^ to check whether Spm promotes the contacts found in this conformation or not. For Tau, clustering of 44 filament structures yielded 14 distinct clusters, with two clusters containing 13 members each and seven unique structures showing no shared contacts. The list of these fibril structures and their corresponding clusters IDs are given in Table S2. The contact maps for these clusters are shown in Fig. S3. For αS, 5 clusters were identified from 7 filament structures (Fig. S4). The list of the 13 αS fibril structures are given in Table S3 along with their corresponding cluster IDs.

Given both diverse disease-relevant fibril structures from experiments and contact maps at different Spm conditions from all-atom simulations, we would like to assess whether the intra-chain contacts observed in the fibril structure are preserved or altered in simulations with varying Spm concentrations. For example, the contact between residue 330 and 359 belonging to Tau-K18 system is shown in a simulation snapshot in Fig. 1E which is observed in numerous fibril structures. For this contact pair, its probabilities at various Spm concentrations has been checked to see whether this contact formation is more or less probable in the presence of Spm. While substantial variability is expected for simulating disordered proteins using all-atom simulations, the slope nevertheless captures whether Spm promotes or disrupts the formation of such contacts. This way, when we check the probabilities for all the contact pairs of the fibril, we get a map similar to the one shown in the bottom right of Fig. 1F. This contact map tells us whether the fibril contacts show increase or decrease in presence of Spm. This same protocol has been applied to different disease causing fibril structures belonging to the three IDPs studied here.

### Divergent Mechanisms of Spermine Modulation in K18, αS, and Aβ

In Fig. 2A to C, we check the Spm’s influence on propensity to form filaments related to the three IDPs. The sequences for these proteins are shown in the left panels. The K18 segment (residues 244-372) shown in Fig. 2A is the microtubule binding region of the longest human tau isoform (441 residues). Four repeat regions within K18 denoted as R1, R2, R3 and R4 (shown in different colours in Fig. 2A) forms the core of the cross structure of tau-paired helical filaments.^54^ The number of positive and negatively charged residues shown with blue and red lines, respectively on the sequence indicate a net positive charge for this system which is likely to create an overall repulsive electrostatic interaction between K18 and Spm. To check how Spm modulates the overall changes in probabilities for the contacts observed in the filament structure belonging to AD, using the cryo-EM structure 5O3L^41^. The fibrillar assembly is shown in the middle panel of Fig. 2A, with one monomer highlighted in a darker shade and zoomed in to illustrate intra-chain interactions. The contact difference map at the bottom with the red and blue dots gives an indication of whether the contact probabilities increase and decrease, respectively, upon Spm interaction. Similar analyses for CBD and Pick’s disease are presented in Figs. S5 and S6, along with the respective filament structures and magnified monomer views.

**Figure 2.**
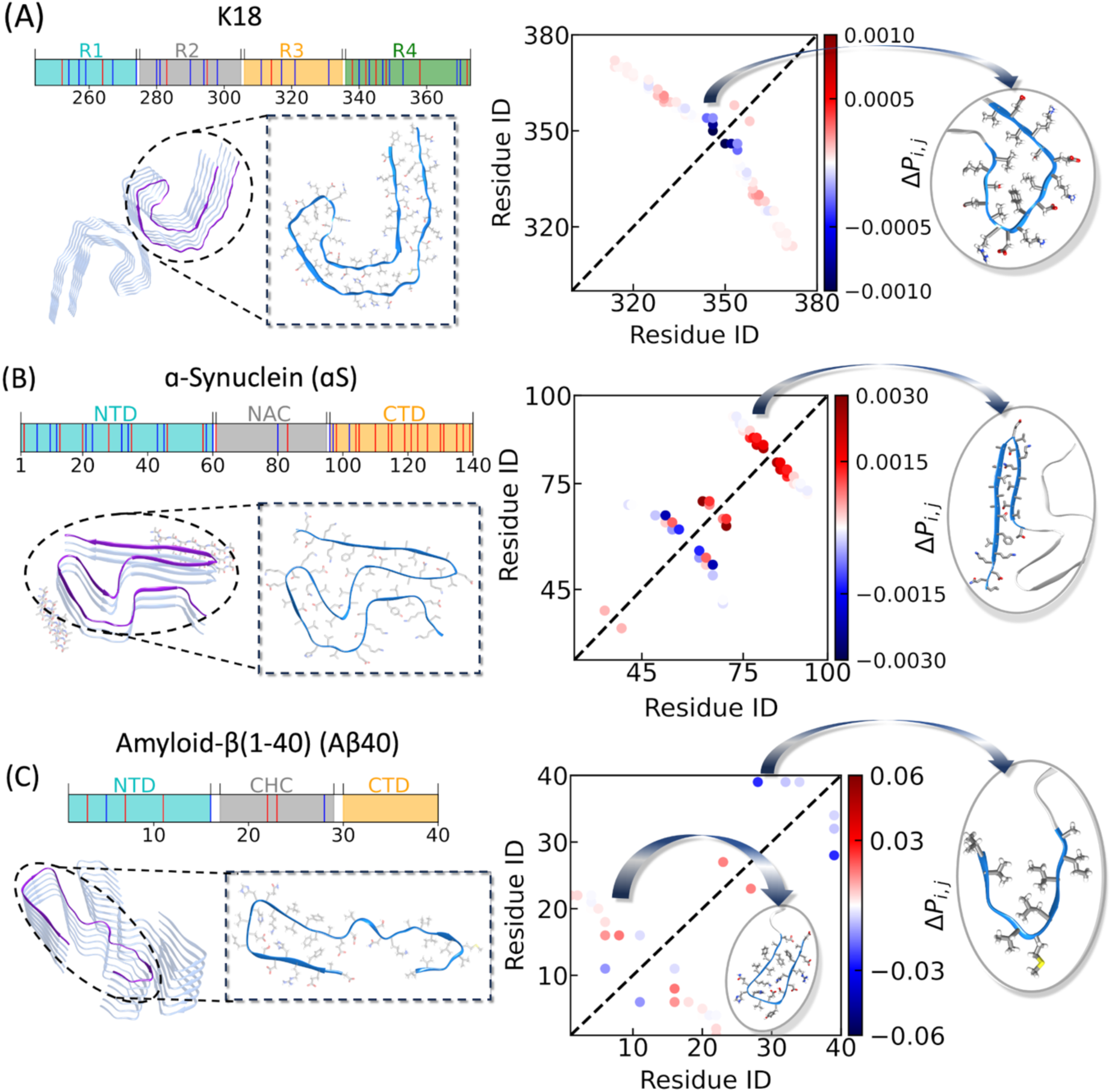
Influence of Spm on the propensities to form disease-relevant native contacts in Tau (A), αS (B), and Aβ40 (C). The top panels on each three subfigures display the sequences of the three intrinsically disordered proteins (IDPs). The middle panels show the corresponding fibril structures—Tau (PDB ID: 5O3L), αS (PDB ID: 8A9L), and Aβ40 (PDB ID: 6SHS)—with one representative monomer highlighted in each case. The bottom panels present the Spm-induced changes in intra-chain contact probabilities for the three systems, with insets highlighting regions that exhibit pronounced alterations in contact formation upon Spm addition.

From the pattern in Fig. 2A, we observe that Spm leads to decrease in fibril-like contacts between residues 340–345, indicating specific binding of Spm at this region. This region corresponds to one end of the C-shaped helical subunit, where charged and hydrophobic residues are densely packed, as illustrated in the inset on the bottom-right of Fig. 2A. This central turn, comprising a heterosteric hydrophobic zipper, has been proposed to form the monomeric hairpin that seeds filament formation in sporadic AD.^36^ Further, this segment has been found to be crucial for governing temperature dependent conformations of Tau.^55^ Since Spm binds near this region and disrupts key intra-chain contacts, it may inhibit the aggregation of Tau into disease-relevant fibrils. For the CBD-related structure (Fig. S5), we also observe a predominance of blue dots, suggesting that Spm interferes with contacts essential for filament formation in this case as well. In contrast, the pattern for Pick’s disease (Fig. S6) is less pronounced, possibly due to the absence of the R2 repeat in this isoform, whereas the Tau sequence used in our simulations includes all four microtubule-binding repeats. Together, these results highlight that Spm engages in specific interactions within the turn region of Tau fibril structure, thereby perturbing a structural element critical for filament nucleation.

The sequence of αS (Fig. 2B) consists of a positively charged CTD, a non-amyloid-β component (NAC) region and a highly negatively NTD as can be seen from the different colours. Presence of several negatively charged residues (red lines) in the sequence are likely to make its interaction with Spm attractive. For this system, we analysed the associated filament structure (PDB ID: 8A9L^48^), shown in the middle panel of Fig. 2B. The intra-chain contact changes, mapped onto the zoomed-in monomer structure, are represented in the contact map at the bottom. In contrast to the Tau filaments, αS exhibits a pronounced increase in contact probabilities, particularly within the NAC region, as highlighted in the inset on the right. A similar trend is observed for the MSA–related filament structure (PDB ID: 6XYO^49^), presented in Fig. S7. Interestingly, despite the structural differences between PD-and MSA-associated αS filaments, the Spm-induced increase in contact probability localizes to the same NAC region in both structures. Previous study on αS and Spm using paramagnetic relaxation enhancement and NMR dipolar couplings indicated that Spm binds primarily to the CTD and weakens its long-range interaction with NTD and NAC region.^56^ This release likely renders the NAC more accessible for intermolecular contacts. Consistent with this, our data suggest that Spm enhances intra-chain contacts within the NAC itself, implying a more global reorganization in contrast to the local structural perturbation observed in Tau^36^. In principle, this shift promotes in general condensation including both aggregation and phase separation. Indeed, in our previous study we demonstrated that Spm extends the lifespan of a *C. elegans* PD model in a concentration-dependent manner by promoting αS LLPS, which facilitates autophagosomes-mediated clearance^57^. This delicate balance between aggregation and LLPS—also observed under varying salt concentrations^58^—highlights a regulatory role of Spm in tuning assembly state of αS.

The Aβ40 sequence shown in Fig. 2C has an overall-3 net charge and therefore is expected to have a net attraction with Spm. The sequence features a CTD, NTD and the central hydrophobic core (CHC); each highlighted with different colours. For Aβ40, we examined the Alzheimer’s disease–associated fibril structure (PDB ID: 6SHS^52^), shown on the middle panel of Fig. 2C with the intra-chain contacts within one monomer zoomed in. The changes in contact probabilities for this system in the absence and presence of Spm is shown in the bottom panel of Fig. 2C. The contact difference map reveals a heterogeneous response: within the CHC and parts of the NTD, Spm enhances several contacts (see insets), while in the CTD and other aggregation-prone regions, Spm reduces contact probabilities. Thus, rather than uniformly stabilizing or destabilizing fibril-like interactions, Spm reshapes the balance of intra-chain contacts across Aβ40 in a region-dependent manner. The experimentally observed increase in Aβ40 aggregation in the presence of Spm^31^ indicates that the promotion of specific contacts outweighs the loss of others, thereby dominating the overall behavior of Aβ40 in solution. A similar behavior is observed for another Aβ40 filament structure (PDB ID: 8QN6^53^) which has similar structural features as the one shown in Fig. 2C indicating consistent yet regionally distinct effects of Spm across Aβ40 filaments.

These results highlight a divergent influence of Spm on different amyloid systems: it promotes filament-like intra-chain contacts in αS (consistent with previous observations^30^), disrupts key nucleation-region contacts in K18, and exerts mixed, region-specific effects in Aβ40. Experimental studies of αS^30^ and Aβ^31^ have already demonstrated Spm’s ability to modulate their aggregation, whereas for the K18 this remains less explored. Our simulations provide a clear, testable hypothesis that Spm should inhibit K18 aggregation, making this construct an ideal system to validate our computational framework. We therefore next assessed the impact of Spm on K18 aggregation using Thioflavin T (ThT) fluorescence assays.

### Experimental Validation of Spermine-Mediated Inhibition of Tau Aggregation

To validate our simulation-based predictions, we monitored the aggregation kinetics of the K18 repeat domain as well as full-length Tau using ThT fluorescence assays in the absence and presence of increasing Spm concentrations (Fig. 3). Aggregation reactions were carried out in 25 mM HEPES buffer (pH 7.4) at 37 °C with agitation, and the ThT signal was recorded over time to capture fibril formation. The kinetic traces were fitted to a sigmoidal function given in Methods section, allowing us to extract phenomenological parameters such as the maximum fluorescence intensity (F_max_) and half-time (t_1/2_) of aggregation. This approach provides a quantitative comparison of how Spm influences fibril growth across different Tau constructs.

**Figure 3.**
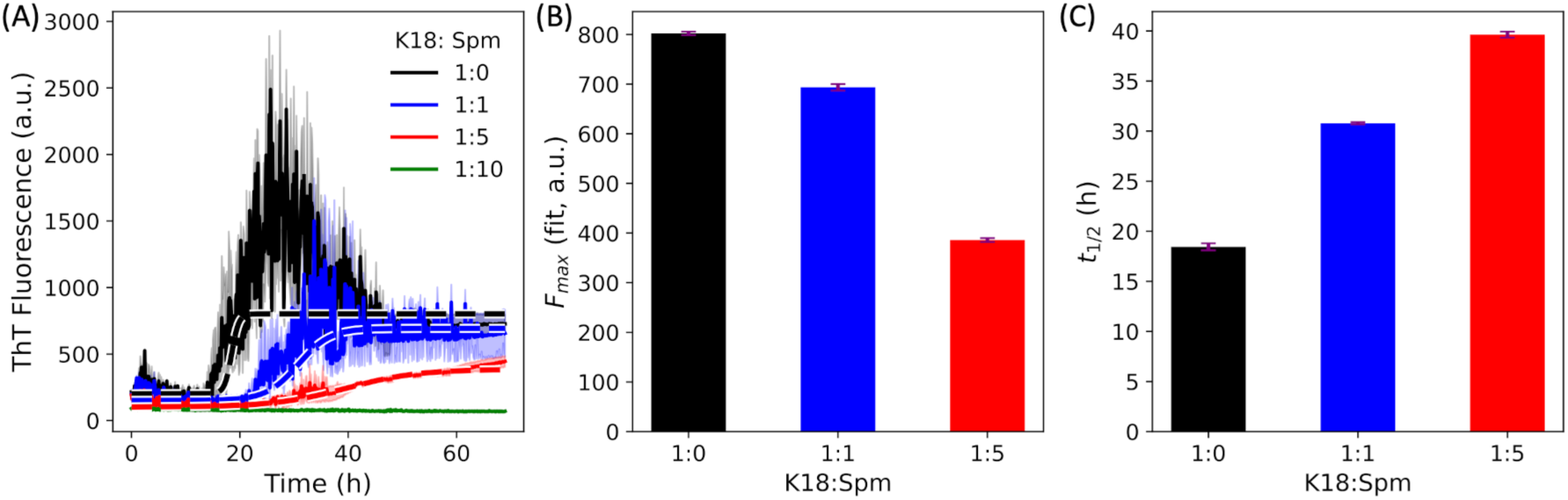
Amyloid aggregation kinetics of K18 monitored by ThT fluorescence in the presence of spermine. ThT fluorescence measurements were performed using 20 μM K18 at varying Spm ratios, as indicated in the legend. (A) Time-dependent ThT fluorescence of K18 in the absence and presence of Spm. Solid lines represent the mean of three independent measurements, with standard error of the mean shown as light-colored shading. Dashed lines indicate sigmoidal fits. For the highest Spm concentration, fitting was not performed. (B, C) Fitted values of the maximum fluorescence intensity (*F*_max_) and aggregation half-time (*t*_1/2_), respectively, at different Spm concentrations. The uncertainty in the fitted parameters was estimated using a bootstrapping procedure.

Fig. 3(A) shows a clear concentration-dependent inhibition of K18 aggregation by Spm. ThT fluorescence intensity progressively decreases with increasing Spm concentration, reaching minimal signal (no aggregates) at the 1:10 K18-Spm ratio, indicating strong suppression of fibril formation. Sigmoidal fits (dashed lines; see Methods) were applied to the 1:0, 1:1, and 1:5 K18:Spm systems, while the 1:10 condition without discernible aggregation was not fitted. The fitted parameters show a systematic reduction in the maximum fluorescence intensity (*F*_max_; Fig. 3B) accompanied by a substantial increase in aggregation half-time (t_1/2_; Fig. 3C), which more than doubles at the 1:5 K18:Spm ratio. These results indicate that Spm markedly slows aggregation kinetics and lowers fibril yield. Consistent with our simulations, this inhibition likely arises from a modest reduction in fibril-prone intramolecular contacts within the K18 monomer, leading to an overall decrease in aggregation propensity.

It is known that truncation of the N-and C-terminal regions of Tau enhances its aggregation propensity.^59^ To assess whether Spm exerts a comparable effect on the aggregation behavior of full-length Tau, parallel experiments using the same protein-to-Spm ratios as for K18 has been carried out on the 441-residue Tau. The fluorescence curves shown in Fig. S8 indicate a much slower aggregation process for the full length Tau in the absence of Spm compared to K18. Such slower aggregation is expected as the N-and C-terminal regions make a fuzzy coat around the aggregation core of Tau. However, in presence of Spm, we see a similar decrease in the fluorescence intensity as observed for K18. This indicates that the subtle changes in the contact formation shown in Fig. 2(A) can also impact the aggregation propensity of full length Tau. These results demonstrate that Spm interferes with the amyloid formation process and further underscore the greater susceptibility of the nucleation-competent K18 construct to Spm-mediated inhibition.

These experiments provide direct validation of our computational framework, establishing K18 as a tractable system in which Spm disrupts aggregation kinetics in agreement with our predictions. Having confirmed this inhibitory effect experimentally, we next turn back to simulations to examine in greater detail how Spm regulates the conformational ensemble of disordered proteins and thereby modulates their aggregation propensities.

### Global Structural Modulation of K18, αS, and Aβ40 by Spm

Having established experimentally that Spm inhibits Tau aggregation, we next returned to simulations to investigate how Spm regulates the conformational ensembles of different amyloidogenic proteins at a global scale. To this end, we first characterized structural features that capture overall compaction and chain organization, including radial distribution functions *g(r)*, the radius of gyration (*R_g_*) and secondary structure preference. These analyses provide a comprehensive view of how Spm reshapes the global conformational behavior of K18, αS, and Aβ, before moving on to residue-specific interactions. Due to the high net charge on Spm, the electrostatic interactions between Spm and proteins are likely to play a pivotal role in governing their behaviour. To probe this, we analysed the radial distribution function (RDF), *g(r)*, which quantifies the spatial distribution between protein Cα atoms and the heavy atoms of Spm for different protein-Spm ratios. The *g(r)* plots for the three systems, presented in Fig. 4A, reveal distinct interaction patterns. As expected for K18 with a net positive charge of +10, the RDFs indicate weak and unfavourable interactions with Spm, evident from the low peak heights in the left panel of Fig. 4A. Conversely, αS with a negative charge of-9 shows strong and favourable interactions with Spm, as seen in the middle panel of Fig. 4A. Notably, the peak heights decrease with increasing Spm concentration, suggesting a saturation of interactions between αS and Spm. Similarly, Aβ40, which also has a net negative charge of-3, exhibits favourable interactions with Spm as can be seen on the right panel of Fig. 4A.

**Figure 4.**
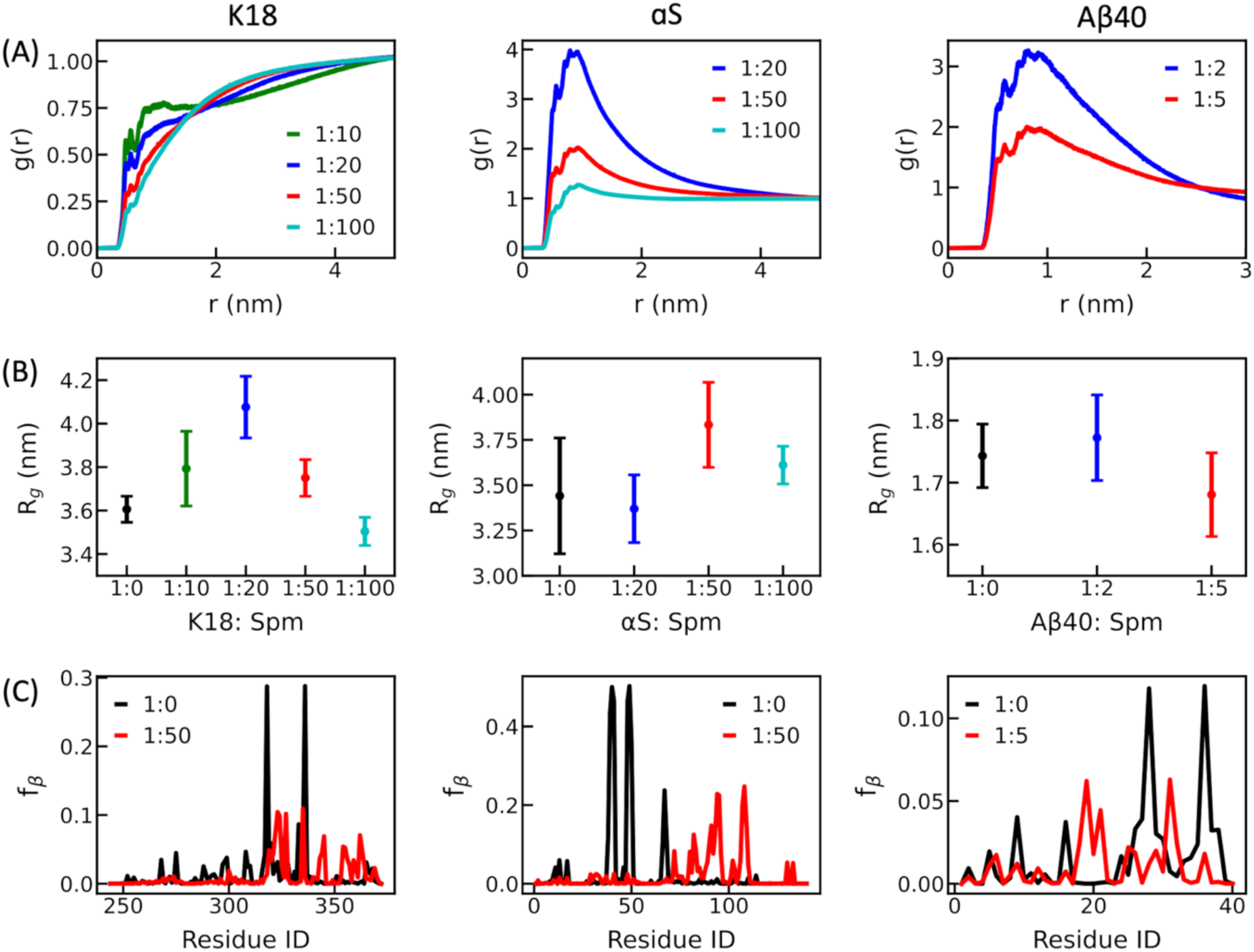
The properties of K18 (left column), αS (middle column) and Aβ40 (right column) in the absence and presence of Spm. (A) The radial distribution function, *g(r)* vs. distances at different Spm concentrations. The *g(r)* has been calculated with respect to the C_α_ atom of all residues from the three systems and the heavy atoms of Spm. (B) The *R_g_* values with respect to Spm ratios for all the systems. (C) The fraction of β-sheet conformation, *f*_β_ for each residue without Spm (black line) and with Spm (red line) for the three systems.

Based on these interaction patterns, one might expect the radius of gyration (*R_g_*) of the protein systems to mirror the trends observed in *g(r)*: attractive association with Spm should neutralize electrostatic repulsion and promote chain compaction. However, the calculated *R_g_* values reveal more complex behavior and in some cases they do not consistently align with the RDF trends, as shown in Fig. 4B. First for K18, the *R_g_* value in the absence of Spm closely matches the experimentally reported value of 3.8 ± 0.3 nm^60^, even though the experimental temperature is slightly higher than the one used here. When Spm is introduced, *R_g_* for K18 initially increases with Spm concentration, as binding to negatively charged residues enhances the net positive charge of K18 and strengthens intrachain electrostatic repulsion. However, at higher Spm concentrations, R_g_ returns to values comparable to the no-Spm condition. Excess Spm screens the electrostatics and limits further charge imbalance, causing *R_g_* to return toward the no-Spm condition. Notably, as K18 expansion has been associated with increased aggregation propensity^61^, these *R_g_* trends imply that Spm-mediated inhibition of Tau aggregation is not a direct consequence of structural compaction induced by Spm binding, but instead point to our earlier observation that localized specific Spm–protein interactions may be at play.

In contrast to K18, for αS, the *R_g_* value without Spm is close to the previous FRET measusrement of 3.3 ± 0.3 nm^62^ and smaller than the previous SAXS measurement of 4.0 ± 0.1 nm^63^, which could be attributed to the temperature and ionic strength differences between the simulation and different experimental setups. With Spm, *R_g_* remains largely unchanged at lower Spm concentrations but increases at higher ratios, reflecting moderate structural expansion. This trend can be explained by the neutralization of negative charges in the CTT, which reduces its long-range attraction to the NTD. Unlike K18, αS does not exhibit a turning behavior, likely because its strong net negative charge prevents saturation within the concentrations tested, so Spm continues to neutralize charges rather than acting as a general electrolyte that primarily provides screening. Meanwhile, Aβ40 exhibits minimal variation in *R_g_* across all Spm concentrations, as shown in the right panel of Fig. 4B. Although Aβ40 demonstrates favourable interactions with Spm, these interactions do not translate into chain expansion. Further insights into these behaviors come from the distributions of *R_g_* values, as shown in Fig. S9. The plot reveals that the *R_g_* distributions for all three systems are broadly similar for K18 and Aβ40, irrespective of Spm presence. However, the distribution shifts slightly to the right at higher Spm ratios for αS. Collectively, these results underscore the complexity of interactions between Spm and these protein systems and highlight the role of specific interactions beyond mere electrostatic effects.

Despite the observed differences in *R_g_* patterns among the three systems, a unifying trend emerges for another structural property, the Flory scaling exponent, *ν*.^64,65^ As shown in Fig. S10, we fitted the root mean squared intrachain distances as a function of the sequence separation *|i-j|* to obtain *ν* for each case using a constant prefactor of 0.55nm as determined from previous literature^66,67^. Table S1 lists ν values for K18, αS, and Aβ40, both with and without Spm. In the absence of Spm, ν values of 0.59, 0.57, and 0.58 for K18, αS, and Aβ40, respectively, indicate that all three systems adopt extended conformations in solution. Upon introducing Spm, *ν* either increases or remains largely unchanged across all systems, irrespective of their net charges, suggesting an overall elongation of the proteins. This observation implies that Spm, despite being positively charged, can interact with the positively charged K18 in a manner that promotes structural elongation.

We next performed secondary structure analysis to check whether a residue on average remains in an helical (H), β-sheet (β) or a random coil (C). Notably, the β-sheet conformation has been implicated in the aggregation behavior of Tau^68^, αS^69^, and Aβ40^70^. Thus, Spm-induced modulation of secondary structure propensities may explain its effects on aggregation. To assess this, we calculated the probabilities of individual residues adopting coil (C), extended (E), and helical (H) conformations in simulations with and without Spm. For all three systems, the fraction of β-sheet conformation (*f*_β_) are shown in Fig. 4C. For clarity, we only show the probabilities for the no-Spm systems and the systems with a protein:Spm ratio of 1:50 for K18 and αS, and 1:5 for Aβ40. The corresponding fractions of coil (*f_C_*) and helical (*f_H_*) conformations are provided in Fig. S11. For K18 (left panel, Fig. 4C), regions with higher extended-state propensity qualitatively agree with previous NMR results^36^. Among these, the 335–340 segment—critical for disease-relevant filament formation as observed in Fig. 2A—shows a notable decrease in *f_E_* upon Spm addition, suggesting a direct influence of Spm on local conformational preferences. For αS (middle panel, Fig. 4C), we observe that in the absence of Spm, residues 36–42 exhibit high *f*_β_, consistent with their known role in aggregation and function^71^. Upon Spm binding, *f*_β_ decreases in this region but increases within the NAC region, where Spm also enhances disease-relevant contacts, pointing to a functional correlation. In Aβ40 (right panel, Fig. 4C), *f*_β_ decreases in the N-terminal domain (NTD) and increases in the central hydrophobic core (CHC) upon Spm addition—mirroring the contact changes observed earlier. These findings underscore the role of extended conformations in promoting aggregation-prone interactions, even at the monomer level.

The three systems highlight distinct behaviors with the presence of Spm: K18 undergoes non-monotonic changes in *R_g_* reflecting initial binding and subsequent saturation and screening; αS displays progressive expansion due to charge neutralization without saturation; and Aβ40 shows limited compaction. These observations suggest that specific amino acid interactions and conformational rearrangements in particular regions, rather than overall net charge and global chain dimensions, play a decisive role in determining structural outcomes. To dissect these effects in greater detail, we next examine changes in secondary structure propensities followed by residue-specific interaction patterns.

### Residue-Specific Interactions Patterns underlying Spm-Mediated Modulation

Finally we would like to explore the specific interactions between Spm and the three systems, as well as the structural and conformational outcomes of these interactions. We start by computing the average number of Spm molecules interacting with each protein residue. This metric, denoted as *N*_Spm_ has been plotted against residue indices for K18, αS and Aβ40 in left, middle and right panel of Fig. 5(A), respectively. The different regions in the sequences are also shown at the top for convinicence. Across all three systems, prominent *N*_Spm_ peaks are observed near negatively charged residues, reflecting the electrostatic nature of Spm–protein interactions. For K18, multiple peaks appear near the start of the R4 repeat, a region enriched in negatively charged residues. This interaction likely explains the observed increase in *R_g_* at lower Spm concentrations. The ^332^PGGG^335^ motif near this site has been found to be crucial for forming the paired helical filaments (PHFs) of Tau.^41^ Binding of Spm near these residues is likely to expand the structure of K18 and is likely to be crucial for preventing the formation of hairpin bend required for fibril assembly. In αS, Spm predominantly interacts with the negatively charged N-terminal region, consistent with its high charge density. While weaker interactions are seen at other negatively charged sites, these are less frequent—aligning with prior NMR chemical shift data for this system^47^. For Aβ40, *N*_Spm_ peaks appear at both termini and the central region, again correlating with the positions of negatively charged residues. The binding patterns observed here suggest that the changes in extended-state conformation seen in Fig. 4C are a direct consequence of Spm interactions. However, the primary binding sites identified in our simulations differ from those showing the largest chemical shift perturbations in earlier NMR studies^31^. To reconcile this, we next examine changes in intra-chain contact maps upon Spm addition, aiming to clarify how Spm binding translates into conformational alterations across these systems.

**Figure 5.**
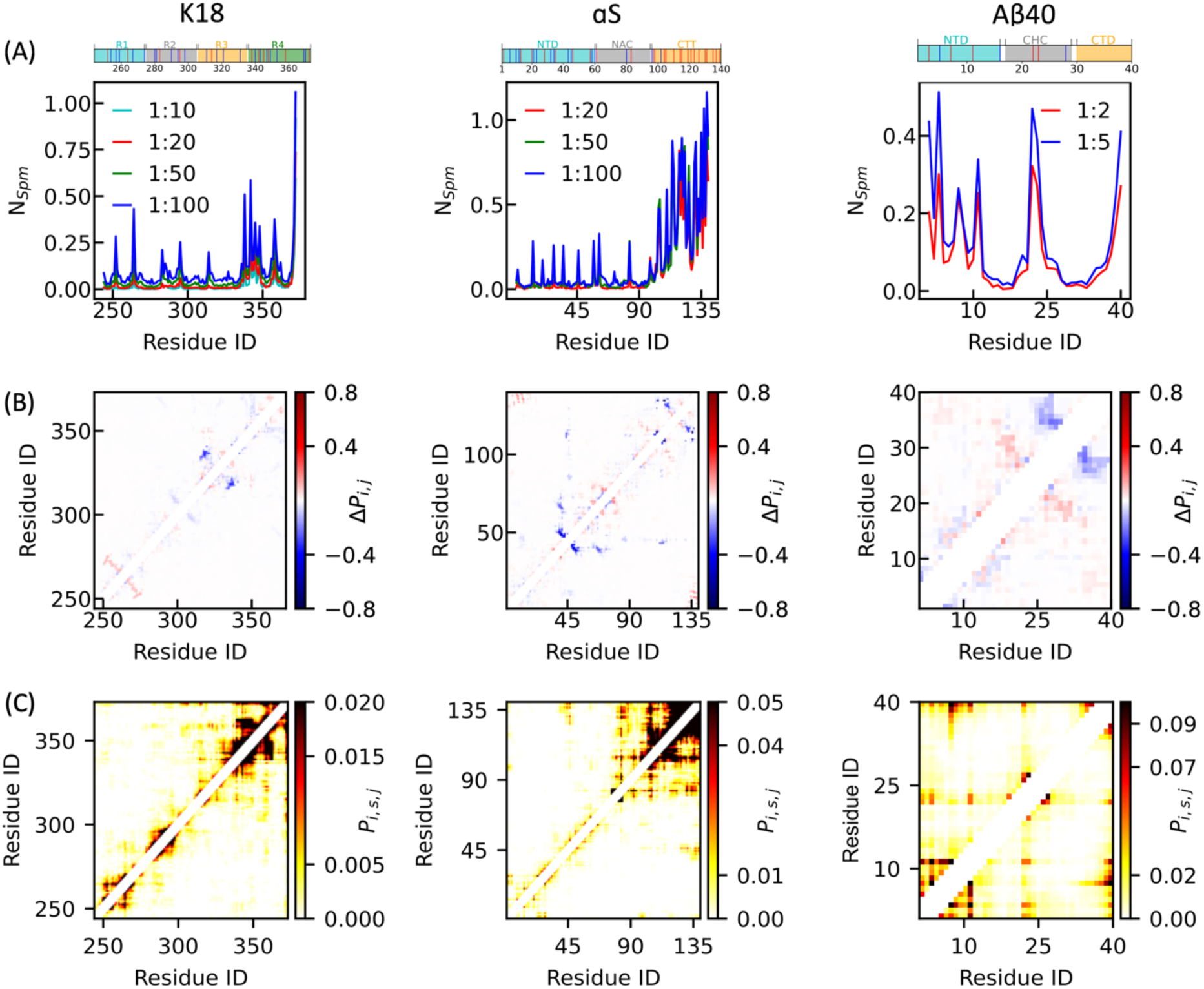
Spermine interactions with K18, αS, and Aβ40 and their effects on intra-residue contacts. (A) Average number of Spm molecules (*N*ₛₚₘ) near each protein residue. (B) Changes in intra-protein contact probabilities (*ΔPᵢ,ⱼ*) for residue pairs, calculated between Spm-free and Spm-bound simulations (1:50 protein-Spm for K18 and αS; 1:5 for Aβ40). (C) Spm-mediated contact probabilities (Pᵢ,ₛ,ⱼ) for residue pairs separated more than three amino acids, measured under the same protein-Spm ratios as in (B).

To investigate how Spm affects intra-chain contacts within these systems, we analysed contacts between heavy atoms of residue pairs separated by at least three residues, both in the presence and absence of Spm. We then computed the difference in contact probabilities by subtracting the values in the Spm-free system from those in the Spm-containing system, indicating whether Spm promotes or disrupts intramolecular contacts. For K18 and αS, the changes in contact probabilities between the Spm-free system and the 1:50 protein–Spm condition are shown in the left and middle panels of Fig. 5B, respectively. For Aβ40, the comparison is between the 1:5 protein–Spm and the Spm-free system, shown in the right panel of Fig. 5B. In K18, the most pronounced reduction in contacts occurs near residues 335–340, coinciding with the region of highest Spm-binding propensity. A smaller decrease is observed near residue 310. Similar behavior has been observed for other Spm ratios also, as shown in Fig. S12. These changes are likely linked to Spm’s ability to modulate Tau aggregation. Prior studies have reported differential interactions between the R1–R4 repeat domains of Tau^20,72–75^, with the R3–R4 segment forming the core of protofibrils in patients with chronic traumatic encephalopathy (CTE)^73^. Spm interaction near the start of the R4 region could therefore influence these aggregation patterns. Notably, the amyloidogenic motif ³⁰⁶VQIVYK³¹¹, located within the R3 repeat, has been shown to drive amyloid formation in vitro^20^ and contribute to pathology in vivo^72^. A reduction in intrachain contacts near this region suggests that the motif may lose its ability to engage upstream sequences, a process known to regulate tau’s aggregation propensity^74^. These findings indicate that, despite the overall electrostatic repulsion between the positively charged K18 and Spm, Spm exhibits selective binding to critical regions in K18, modulating its intramolecular contacts and potentially influencing its aggregation into disease-relevant fibrils.

For αS, Spm shows change in contacts at several places with the most notable decrease in contacts at NTD and between NTD and CTD. Additionally, slight increase in contacts have also been observed at the NAC region across all the Spm ratios as can be seen in middle panel of Fig. 5B and in Fig. S13. These changes resembles the effects seen upon truncation of the CTD, ^76,77^ suggesting that Spm coats this region and thereby exposes the NAC domain, making it more susceptible to intra-and inter-chain interactions. ^77,78^Another notable contact change occurs near residue 46, a site implicated in fibril-like contact formation. The E46K mutation, which introduces a positive charge at this position, is known to enhance αS fibrillization.^78^ Since Spm is likely to bind at or near residue 46 and similarly alter local electrostatics, it may promote aggregation through a mechanism akin to that of the E46K mutation.

For Aβ40, the contact maps indicates a mixed pattern of changes upon Spm binding, as seen in the right panel of Fig. 5B and in Fig. S14. Notably, the CHC and CTD regions show distinct responses: intramolecular interactions within the CHC increase, whereas those within the CTD decrease. In addition, enhanced interactions are observed within CHC, particularly between residues 16-20 and 27-30. This effect likely arises from Spm binding to residues ^22^Glu-^23^Asp (Fig. 5A), which lie between the two fragments. By neutralizing negative charges, Spm reduces electrostatic repulsion within this region. The two fragments showing significant contact variations upon Spm addition align with previous NMR study on Aβ40 with Spm, where residues with the largest chemical shift changes corresponding to positions near residues 4-5, 15-17 and 27-28^31^. Similar effects have also been documented in the presence of positively charged metal ions ^79^ and other small molecules^80^.

In addition to analysing changes in contact maps in the presence of Spm, we also examined Spm-mediated intra-chain contacts in these systems. Fig. 5C presents the Spm-mediated contact maps for K18 (left), αS (middle), and Aβ40 (right). For K18, the plot reveals that Spm-mediated contacts are predominantly short-range. Although a major change in contact probability is observed near residues 335–340, consistent with the region of strongest Spm binding, several upstream residues also display Spm-mediated interactions. In the case of αS, the most prominent Spm-mediated contacts occur within the CTD, consistent with the higher Spm binding observed in this region. Interestingly, we also detect long-range contacts between the NTD and CTD, likely facilitated by Spm. This interaction may disrupt normal interactions between the NAC region and NTD, thereby increasing the aggregation propensity of αS by freeing the NAC domain for self-association. For Aβ40, the most probable Spm-mediated interactions are found within the NTD and between the NTD and CTD. Although residues 23–24 interacts with Spm, the right side plot in Fig. 5C clearly indicate that the interaction between the regions of NTD and CHC observed in Fig. 5B does not come from Spm mediation but direct residue-residue interactions.

Together, these results demonstrate that Spm can facilitate both short-range and long-range interactions within IDPs. In K18, the distribution of negatively charged residues drives Spm to bind regions critical for aggregation, potentially disrupting key contacts. In αS, Spm binding at multiple acidic sites reduces the CTT-NAC interactions and increases solvent accessibility of the aggregation-prone NAC. For Aβ40, aggregation of the central hydrophobic core (CHC) is normally hindered by surrounding acidic residues; however, Spm binding likely neutralizes electrostatic repulsion, enabling CHC-driven aggregation. These findings suggest that Spm modulates the aggregation of these three IDPs via distinct mechanistic pathways, each shaped by the protein’s charge distribution and structural organization, underscoring the need to consider sequence-and region-specific effects when studying polyamine–protein interactions.

## Discussion

Despite significant advances in experimental and computational studies, unravelling the drivers of IDP aggregation and identifying strategies to prevent it remains a formidable challenge. One promising avenue involves examining the influence of cellular components, which may indirectly modulate the formation of disease-relevant assemblies. Among these, polyamines represent a class of small, positively charged molecules that have been implicated in tauopathies and synucleinopathies. In this study, we investigate the impact of one such polyamine, spermine, on the aggregation behaviors of three representative IDPs: the K18 region of Tau, full length αS, and Aβ40. Using long-timescale all-atom explicit-solvent molecular dynamics simulations and ThT fluorescence assays, we probe how Spm affects the early stages of disease-relevant filament formation. Given growing evidence that small oligomers may be toxic^81^, and that aggregation may be initiated at the monomeric level^36^, monomeric all-atom simulations provide a valuable lens to identify residue-specific interactions that shape the aggregation process.

To probe how Spm influences fibril-like interactions, we examined intra-chain native contact propensities derived from fibril structures. Our findings show a reduction in contact probabilities for Tau filament structures associated with Alzheimer’s and Pick’s disease in the presence of Spm. Conversely, αS filament contacts linked to Parkinson’s and multiple system atrophy (MSA) exhibited increased probabilities. Aβ40 displayed a mixed response. These observations were supported by ThT fluorescence experiments for K18, which showed decreased aggregation in the presence of Spm. For αS, the enhancement of aggregation by Spm is consistent with prior findings. However, the ability of Spm to also promote liquid-like condensate formation for αS under crowding condition also plays a crucial role in controlling the progress of different diseases, as was observed in our previous study^37^. The fate of aggregation and LLPS of αS may depend on a combination of different factors such as mutations, salt or cellular polyamines.

To further elucidate the conformational influence of Spm, we analysed both global and local structural changes. The net attraction or repulsion observed in the radial distribution functions could be fully explained by the overall net charge of each protein chain. However, radius of gyration values did not follow a simple monotonic trend with Spm concentration, indicating specific Spm–IDP interactions beyond global charge effects. Secondary structure analysis further revealed notable changes at regions known to be important for fibril formation. Residue-specific binding analysis demonstrated preferential interactions of Spm with specific segments in each IDP, with clear structural consequences: in K18, the region with the strongest Spm binding coincided with major contact disruptions in the aggregation core; in αS, Spm primarily bound to the C-terminal domain (CTD), reducing CTD–NAC interactions and altering NAC accessibility; and in Aβ40, binding near residues 23–24 was accompanied by local compaction of adjacent regions. Thus, Spm exerts divergent, protein-specific influences on aggregation.

The lack of well-defined binding pockets in IDPs presents a major obstacle in traditional drug design. As such, exploring alternative strategies, such as modulating cellular polyamine levels, may offer novel therapeutic avenues for controlling IDP aggregation. Our findings suggest that regulating spermine concentration could potentially modulate IDP aggregation, and more broadly reshape the conformational landscapes of disease-associated IDPs. Importantly, all-atom simulations of monomeric proteins prove valuable in revealing residue-specific interactions that underlie these effects, offering mechanistic insight into how small molecules like polyamines modulate complex aggregation behaviors. Future work should extend these approaches to more aggregation-prone variants and cellular contexts to evaluate the therapeutic potential of targeting polyamine–protein interactions.

## Methods

### All-atom MD simulation

The all-atom MD simulations were performed using GROMACS version 2023.3 software package^82,83^ for all the systems. We have used AMBER ff99SBws^35^ force field for the proteins along with TIP4P2005 water model^84^, and general AMBER force field^85^ (GAFF) for Spm. The initial structures for all the three systems were inserted in a triclinic dodecahedron simulation box. Following this, required number of Spm molecules were added. The simulation boxes are filled with water molecules and required number of sodium and chloride ions are added to obtain ∼150 mM NaCl concentration along with ions to neutralize the systems. Antechamber module^86^ has been used to generate the GAFF parameters for Spm. Atomic partial charges were derived using the restrained electrostatic potential (RESP) fitting procedure^87^ based on electrostatic potentials computed at the HF/6-31G* level of theory. The systems were first energy minimized using steepest descent method. This was followed by a 1ns equilibration simulation under constant pressure and temperature using Berendsen pressure coupling^88^ and V-rescale temperature coupling^89^. The same coupling constant of 1ps has been used for both temperature and pressure coupling during the equilibration. For the K18 systems, the simulations were run at 283 K in accordance to the NMR experiments carried out for this system. For αS and Aβ40, the simulations were carried out at 288 and 278 K, respectively to make the simulation conditions consistent with previously reported NMR experiments on these systems.^31,90^ equilibrated structures were used for production simulation under constant pressure and temperature condition. Parrinello-Rahman barostat^91^ has been used along with V-rescale thermostat for these simulations with a coupling constant of 1ps for both barostat and thermostat. Electrostatic interactions were treated using PME electrostatics^92^ with 1.2 nm cutoff. van der Waals cutoff was set to 1.2 nm. Initial 500ns has been discarded from K18 and Aβ40 simulations while for αS simulations, initial 200ns has been discarded. We calculated the percentage of trajectory frames in which the minimum distance to the periodic image was less than 0.3 nm and found it to be negligible (Table S1). All subsequent analyses were performed after excluding these frames.

### Simulation analysis

The radial distribution functions, *R_g_* values and secondary structure assignments were obtained using GROMACS for all the systems. The *R_g_* values are given in Table S1, and the errors were estimated using a block averaging method with four blocks. The DSSP module^93^ of GROMACS has been used to assign the secondary structure for each residue of the proteins. From these assignments, the probability of each residue in coil, helix or sheet conformation has been determined. For calculating the number of Spm around each protein residue, number of Spm that comes in contact with each residues have been calculated. For this purpose, a contact has been considered between Spm and residues if any of their non-hydrogen atoms come within a distance of 0.45 nm of each other. Same condition has been used for obtaining the intra-residue contact maps for all the systems. In all the cases, residues that are separated by at least three other residues have been considered. For Spm mediated contacts, the distance cutoff chosen between non-hydrogen atoms of Spm and protein residues is 0.6nm. In this case also, we have taken only those residues that are separated by at least three other residues.

### Protein purification

Full-length human Tau (441 amino acids, UniProt P10636-8) and K18 variant (residues 244-372) were expressed in *E. coli* BL21 (DE3) and purified as previously described^94^. Expression was performed at 37°C in LB medium with ampicillin (100 mg/L). IPTG (0.4 mM final concentration) was added at OD600 0.6-0.8, followed by 2 h cultivation. Cells were harvested and stored at-20°C. Cell pellets were resuspended in 50 mM NaPi, 2.5 mM EDTA, pH 6.2 with protease inhibitors. Soluble extracts were obtained by sonication and centrifugation. Supernatants were heated at 75°C for 15 min, then centrifuged. Proteins were purified using Hi-Trap SP FF cation exchange chromatography with NaCl gradient elution (50 mM NaPi, 2.5 mM EDTA, 500 mM NaCl, pH 6.2). Purified proteins were verified by 12% SDS-PAGE, dialyzed against 20 mM ammonium bicarbonate, and lyophilized. Concentrations were determined by UV absorption at 280 nm (extinction coefficients: Tau 7450 M⁻¹ cm⁻¹, K18 1490 M⁻¹ cm⁻¹).

### ThT kinetics

20 μM Tau and K18 were incubated with varying spermine ratios in the presence of two glass beads (1.0 mm diameter, Sigma-Aldrich) and 50 μM ThT in 25 mM HEPES (pH 7.4). Samples were prepared on ice, and 20 μL aliquots were dispensed into 384-well black/clear-bottom microplates (ThermoFisher, cat. no. 242764) and sealed using aluminum adhesive foil (neoLab). Aggregation was monitored using a PHERAstar FSX microplate reader (BMG LABTECH, Germany) at 37°C with 300 rpm shaking. ThT fluorescence was measured every 7 minutes (excitation 430 nm, emission 480 nm). Each condition was measured in three repeats. The mean and the standard error of the mean were calculated for subsequent analyses. The ThT fluorescence time-course data were fitted to a sigmoidal growth model using the logistic function:

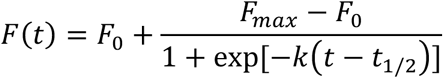

where *F(t)* is the fluorescence intensity at time *t*. *F_0_* represents the initial baseline fluorescence prior to aggregation, and *F_max_* denotes the final plateau fluorescence after aggregation reaches completion. The parameter *k* is the apparent rate constant describing the steepness of the growth phase, with larger values indicating a faster transition. The term *t_1/2_* corresponds to the half-time of the reaction, defined as the time at which fluorescence reaches halfway between *F_0_* and *F*_max_. Lag time was calculated as *t_1/2_-2/k*, consistent with the standard tangent-intersection method used for sigmoidal ThT kinetics. The uncertainty in the fitted parameters was estimated using a bootstrapping procedure.

## Acknowledgement

The authors acknowledge the support from the National Institutes of Health (R35GM146814, W.Z.), Swiss National Scientific Foundation (10002967, J.L.), and the Research Computing at Arizona State University.

## SUPPORTING INFORMATION

## Supporting Tables

**Table S1.** Simulation details for all the systems along with their radius of gyration (*R_g_*) and Flory scaling exponent (ν) values.

| System | Protein-Spm ratio | Box size (nm) | No. of atoms | Simulation lengths (μs) | % of Frames (Min. dist. ≥ 0.3 nm) | $R_g$ (nm) | $\nu$ |
| --- | --- | --- | --- | --- | --- | --- | --- |
| K18 | 1:0 | $12.08 \times 12.08 \times 8.54$ | 166881 | 5 | 95.6 | 3.61 | 0.594 |
| | 1:10 | $12.09 \times 12.09 \times 8.55$ | 168233 | 5 | 97.6 | 3.79 | 0.600 |
| | 1:20 | $12.07 \times 12.07 \times 8.53$ | 166553 | 5 | 94.1 | 4.08 | 0.621 |
| | 1:50 | $12.07 \times 12.07 \times 8.53$ | 167513 | 5 | 95.9 | 3.75 | 0.598 |
| | 1:100 | $12.12 \times 12.12 \times 8.57$ | 169113 | 5 | 99.4 | 3.50 | 0.589 |
| αS | 1:0 | $14.89 \times 14.89 \times 10.53$ | 312963 | 2 | 100.0 | 3.44 | 0.566 |
| | 1:20 | $14.97 \times 14.97 \times 10.59$ | 316691 | 2 | 100.0 | 3.37 | 0.577 |
| | 1:50 | $14.93 \times 14.93 \times 10.56$ | 314563 | 2 | 99.4 | 3.83 | 0.590 |
| | 1:100 | $14.97 \times 14.97 \times 10.59$ | 317387 | 2 | 100.0 | 3.61 | 0.573 |
| Aβ40 | 1:0 | $7.17 \times 7.17 \times 5.07$ | 34911 | 5 | 99.8 | 1.74 | 0.580 |
| | 1:2 | $7.17 \times 7.17 \times 5.07$ | 34931 | 5 | 96.6 | 1.77 | 0.594 |
| | 1:5 | $7.17 \times 7.17 \times 5.07$ | 34979 | 5 | 99.8 | 1.68 | 0.570 |

**Table S2.** List of PDBs for Tau fibrils belonging to different clusters:

| Cluster Id | PDB file name |
| --- | --- |
| Cluster 1 | 7QJV<sup>1</sup>, 7QL4<sup>1</sup>, 7QKI<sup>1</sup>, 7QKK<sup>1</sup>, 5O3L<sup>2</sup>, 5O3T<sup>2</sup>, 5O3O<sup>2</sup>, 6HRE<sup>3</sup>, 7NRS<sup>4</sup>, 7NRX<sup>4</sup>, 7NRQ<sup>4</sup>, 7NRV<sup>4</sup>, 7NRT<sup>4</sup> |
| Cluster 2 | 7QJW<sup>1</sup>, 7QJZ<sup>1</sup>, 7QK5<sup>1</sup>, 7QKU<sup>1</sup>, 7QKV<sup>1</sup>, 7QKX<sup>1</sup>, 7QL0<sup>1</sup>, 7QL1<sup>1</sup>, 7QKG<sup>1</sup>, 7QK6<sup>1</sup>, 7QKW<sup>1</sup>, 6NWP<sup>5</sup>, 7QK1 (PF1*)<sup>1</sup> |
| Cluster 3 | 7QL3<sup>1</sup>, 7QJY<sup>1</sup>, 7QKJ<sup>1</sup>, 7QK2<sup>1</sup>, 7QK1 (PF2*)<sup>1</sup> |
| Cluster 4 | 7QKL<sup>1</sup>, 7R4T<sup>1</sup>, 7QJX<sup>1</sup> |
| Cluster 5 | 6TJX<sup>6</sup>, 6TJO<sup>6</sup> |
| Cluster 6 | 7R5H<sup>1</sup> |
| Cluster 7 | 7QKF<sup>1</sup> |
| Cluster 8 | 7QL2<sup>1</sup> |
| Cluster 9 | 7QKZ<sup>1</sup> |
| Cluster 10 | 7QK3<sup>1</sup> |
| Cluster 11 | 7QKH<sup>1</sup> |
| Cluster 12 | 7QKY<sup>1</sup> |
| Cluster 13 | 7QKM<sup>1</sup> |
| Cluster 14 | 6GX5<sup>7</sup> |

\*PF1 and PF2 indicate two conformationally different protofilaments belonging to the same PDB entry.

**Table S3.** List of PDBs for αS fibrils belonging to different clusters.

| Cluster Id | PDB file name |
| --- | --- |
| Cluster 1 | 6SST<sup>8</sup>, 6SSX<sup>8</sup> |
| Cluster 2 | 6XYP<sup>9</sup>, 6XYQ<sup>9</sup> |
| Cluster 3 | 6H6B<sup>10</sup> |
| Cluster 4 | 8A9L<sup>11</sup> |
| Cluster 5 | 6XYO<sup>9</sup> |

**Table S4.**
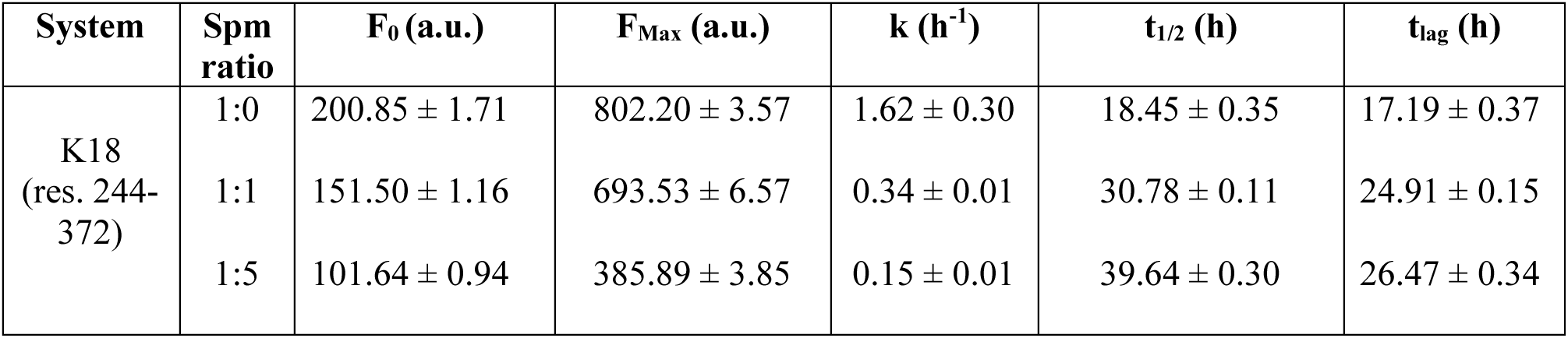
Phenomenological and fitting parameters for amyloid aggregation for K18, derived from sigmoidal curve fitting to Thioflavin T (ThT) fluorescence kinetics data.

| System | Spm ratio | F<sub>0</sub> (a.u.) | F<sub>Max</sub> (a.u.) | k (h<sup>-1</sup>) | t<sub>1/2</sub> (h) | t<sub>lag</sub> (h) |
| --- | --- | --- | --- | --- | --- | --- |
| K18<br>(res. 244-372) | 1:0 | 200.85 ± 1.71 | 802.20 ± 3.57 | 1.62 ± 0.30 | 18.45 ± 0.35 | 17.19 ± 0.37 |
|  | 1:1 | 151.50 ± 1.16 | 693.53 ± 6.57 | 0.34 ± 0.01 | 30.78 ± 0.11 | 24.91 ± 0.15 |
|  | 1:5 | 101.64 ± 0.94 | 385.89 ± 3.85 | 0.15 ± 0.01 | 39.64 ± 0.30 | 26.47 ± 0.34 |

## Supporting Figures

**Figure S1.**
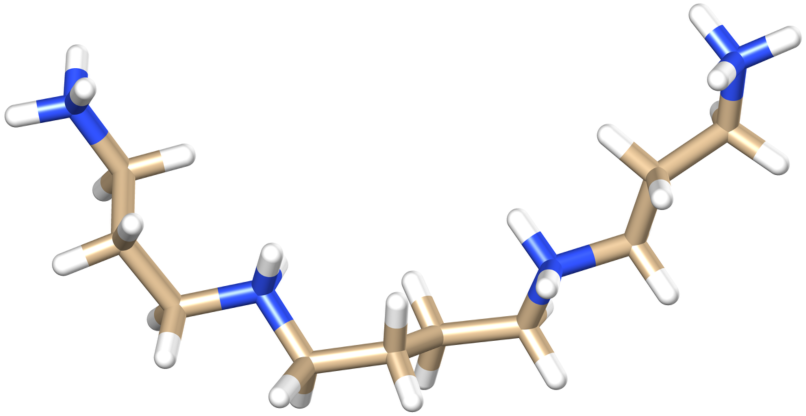
The structure of spermine (Spm) with a net charge of +4.

**Figure S2.**
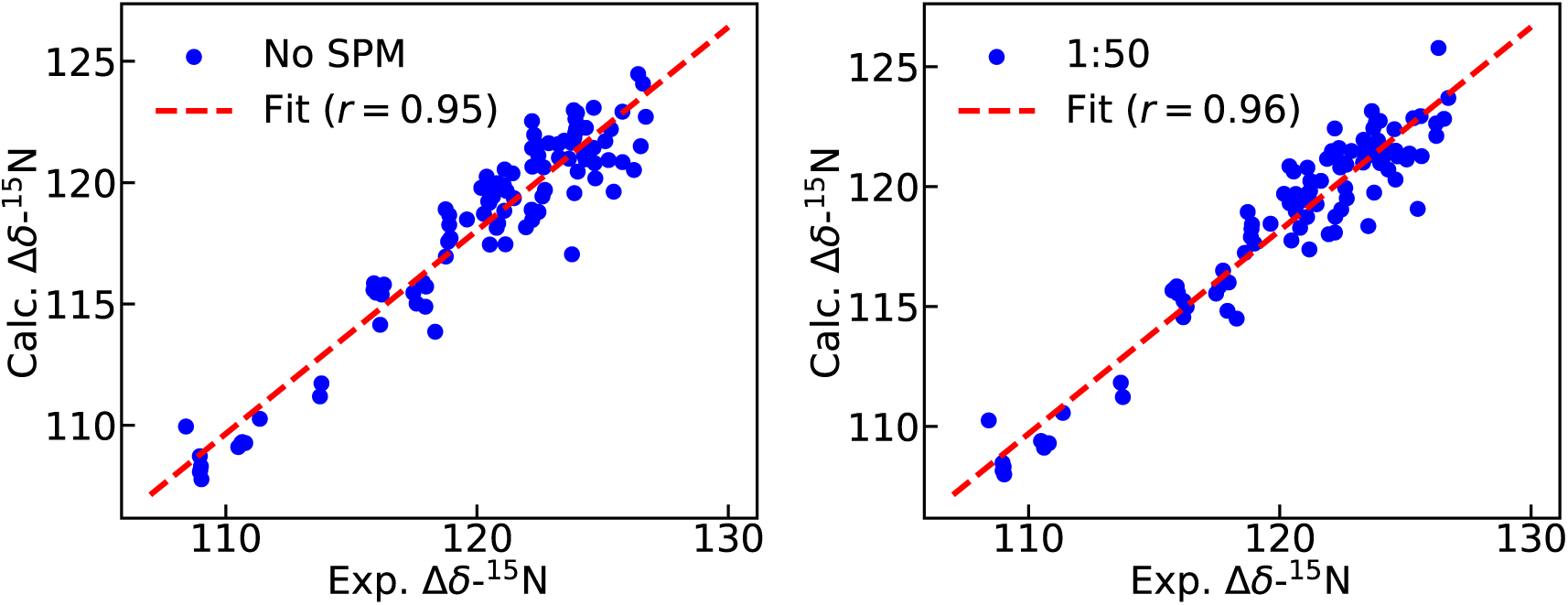
Comparison between experimental and calculated NMR chemical shifts. Calculated vs experimental NMR chemical shifts for ^15^N atoms of K18 system in the absence (left) and presence of Spm (right) at 1:50 protein-Spm ratio. The red dashed line shows the linear fitting of the data with Pearson correlation coefficient (*r*) values given in the plots.

**Figure S3.**
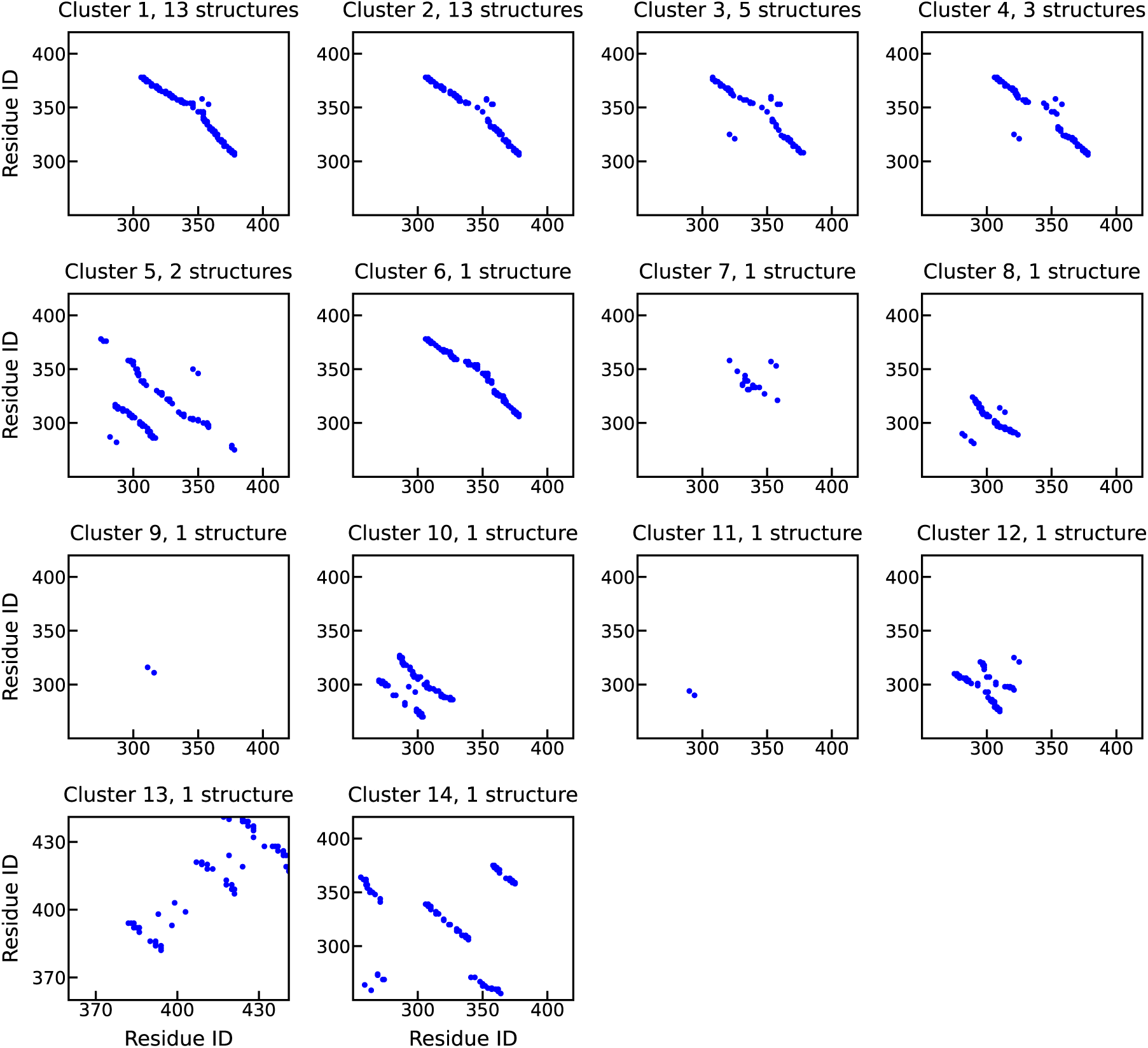
Contact map of different Tau fibril structures. Intra-residue contacts maps of the 14 clusters obtained from Tau fibrils. The number of structures belonging to each cluster is mentioned at the top of each contact map.

**Figure S4.**
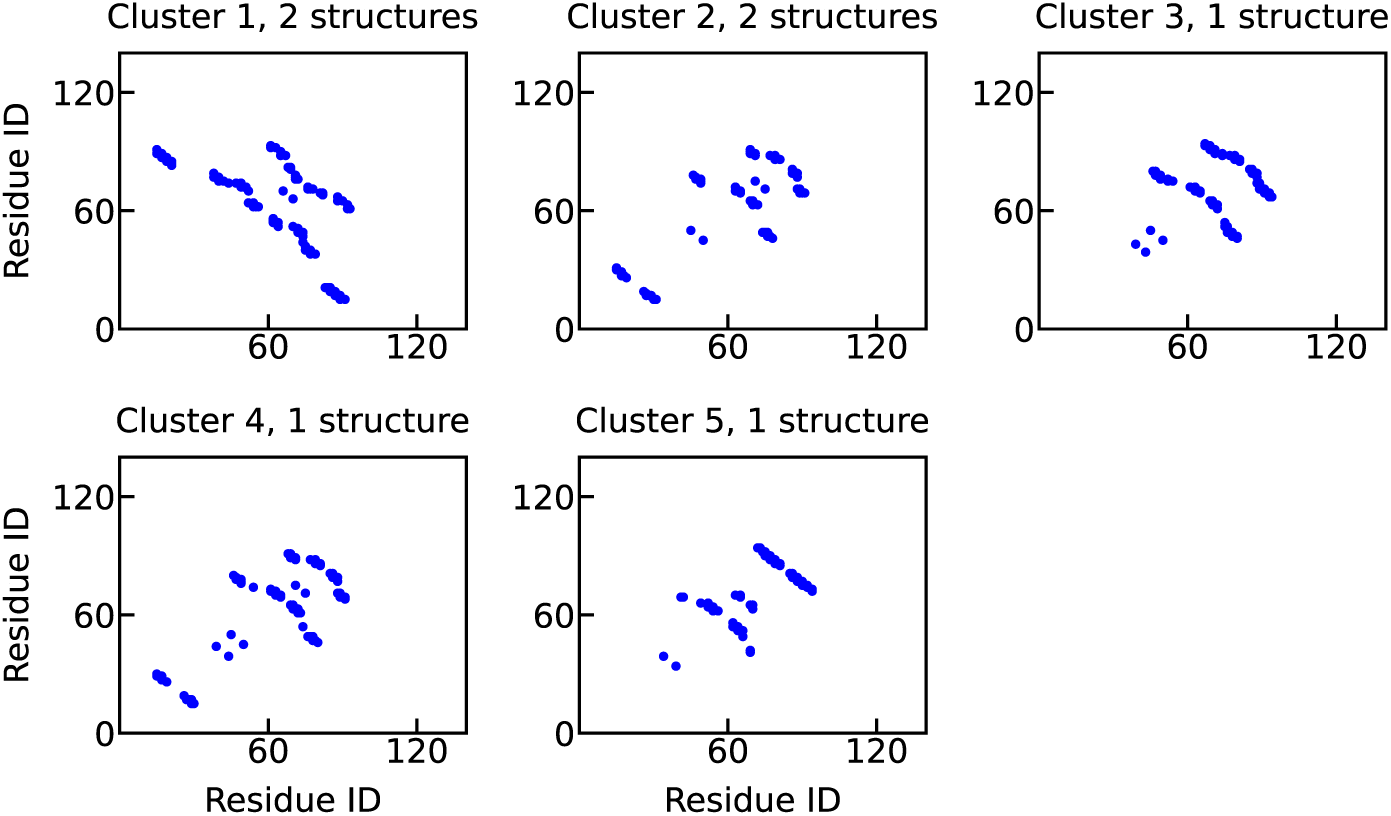
Contact map of different. αS **fibril structures.** Intra-residue contact maps of the five clusters obtained from αS fibrils. The number of structures belonging to each cluster is mentioned at the top of each contact map.

**Figure S5.**
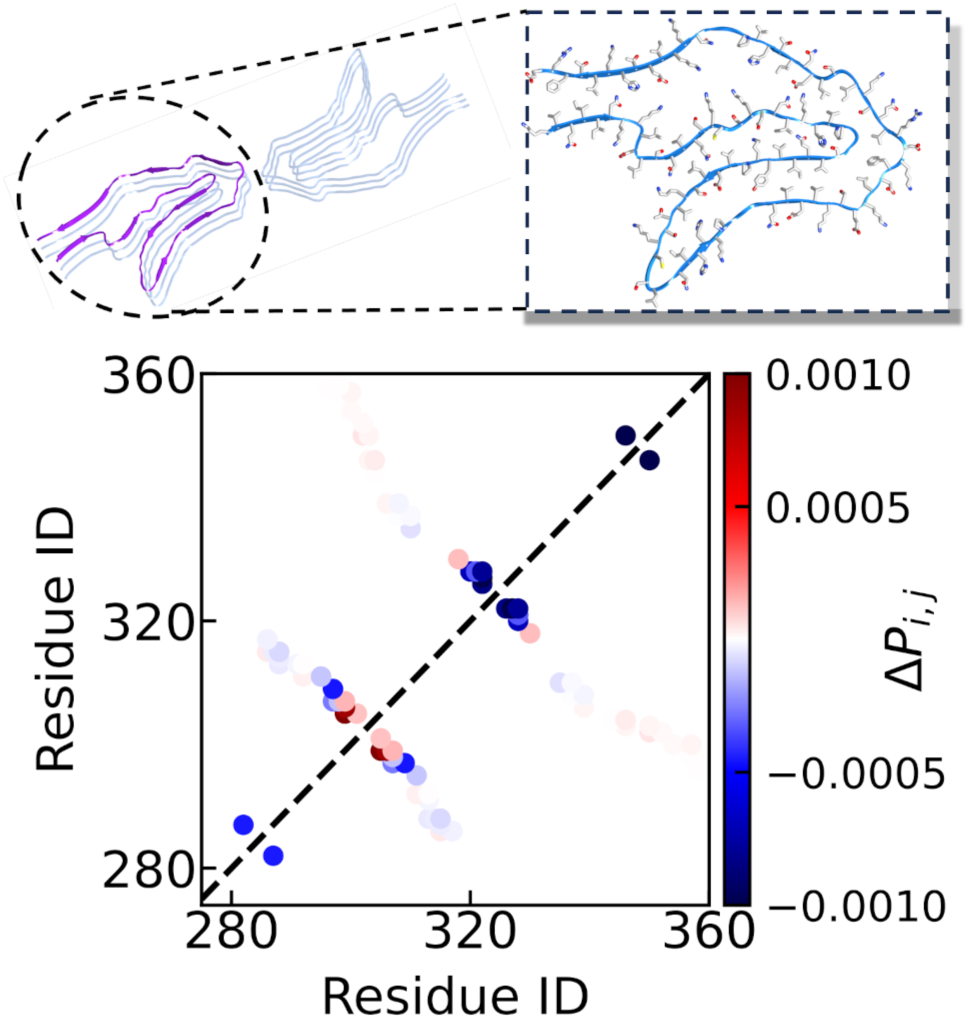
Structure and change in contact probabilities for Tau fibril associated with Corticobasal Degeneration (CBD). The fibril structure of CBD related Tau fibril (PDB ID: 6TJX) with one monomer highlighted in purple and shown in the zoomed in structure on the right. The corresponding contact map showing the changes in probabilities in presence of Spm for contact pairs present in the fibril structure.

**Figure S6.**
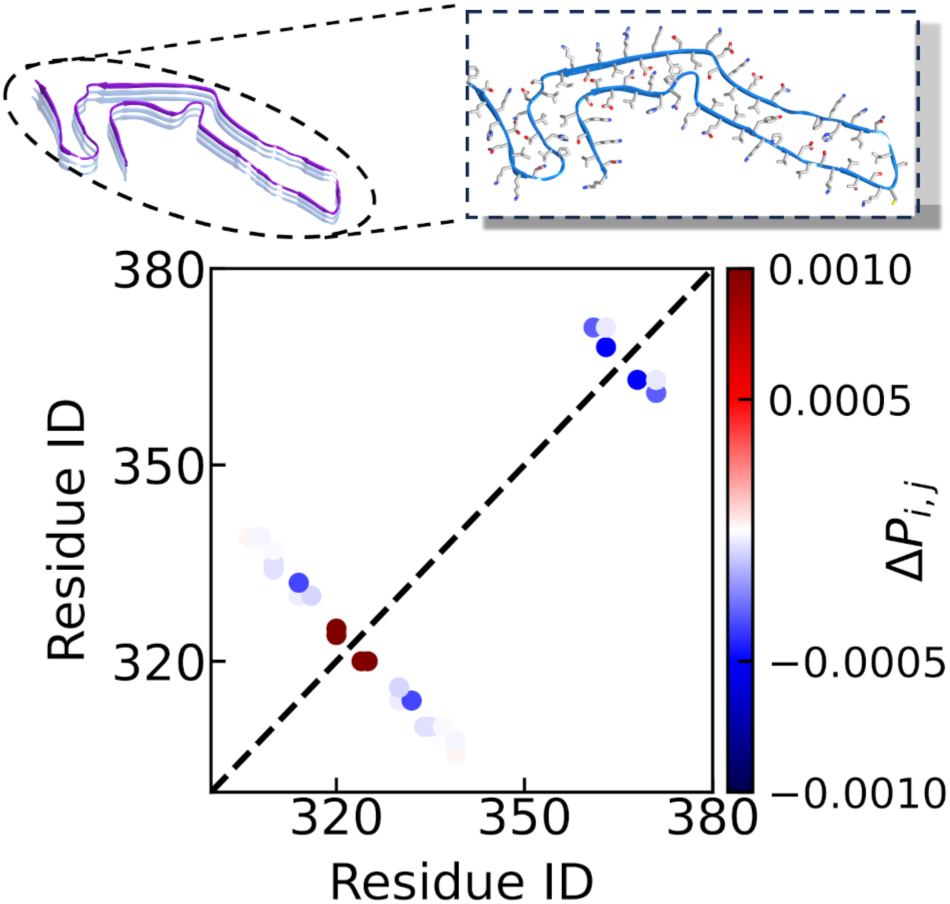
Structure and change in contact probabilities for tau fibril associated with Pick’s disease. The fibril structure of Pick’s disease related tau fibril (PDB ID: 6GX5) with one monomer highlighted in purple and shown in the zoomed in structure on the right. The corresponding contact map showing the changes in probabilities in presence of Spm for contact pairs present in the fibril structure.

**Figure S7.**
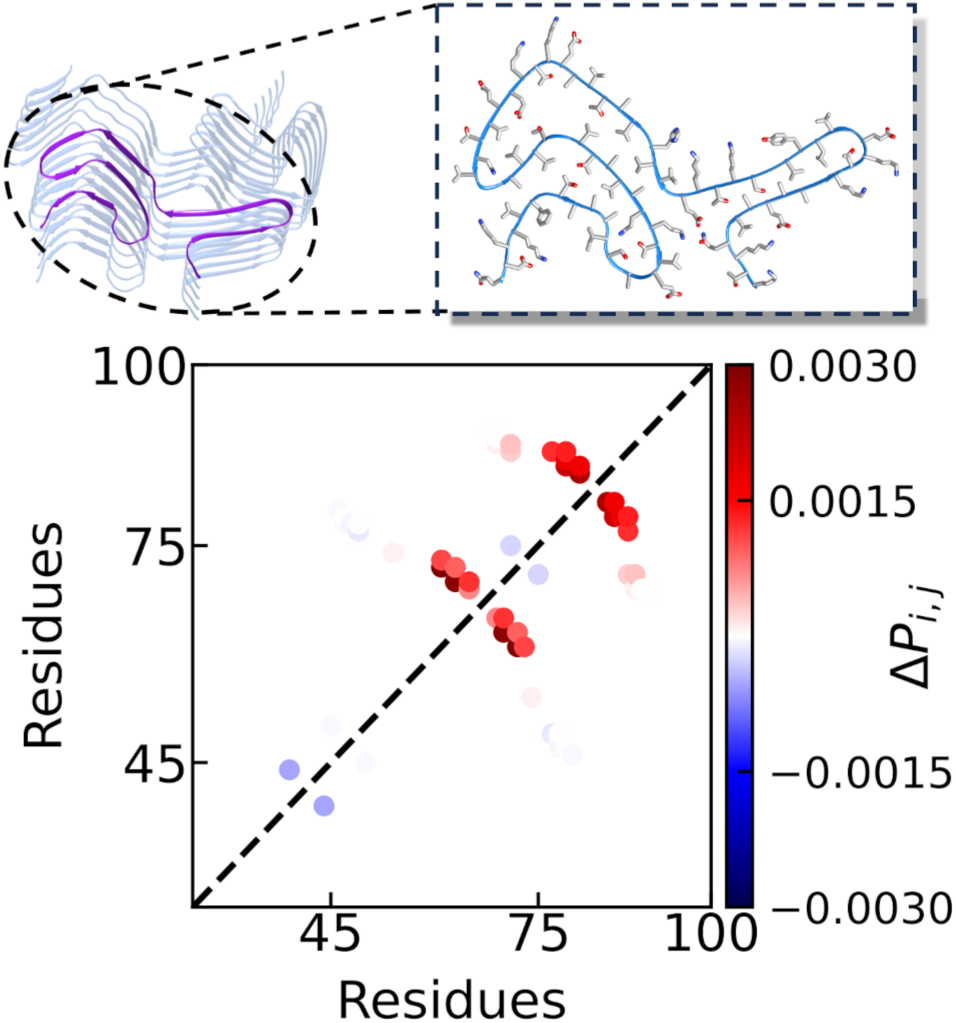
Structure and change in contact probabilities for αS fibril associated with Multiple System Atrophy (MSA). The fibril structure of MSA related αS fibril with one monomer highlighted in purple and shown in the zoomed in structure on the right. The corresponding contact map showing the changes in probabilities in presence of Spm for contact pairs present in the fibril structure.

**Figure S8.**
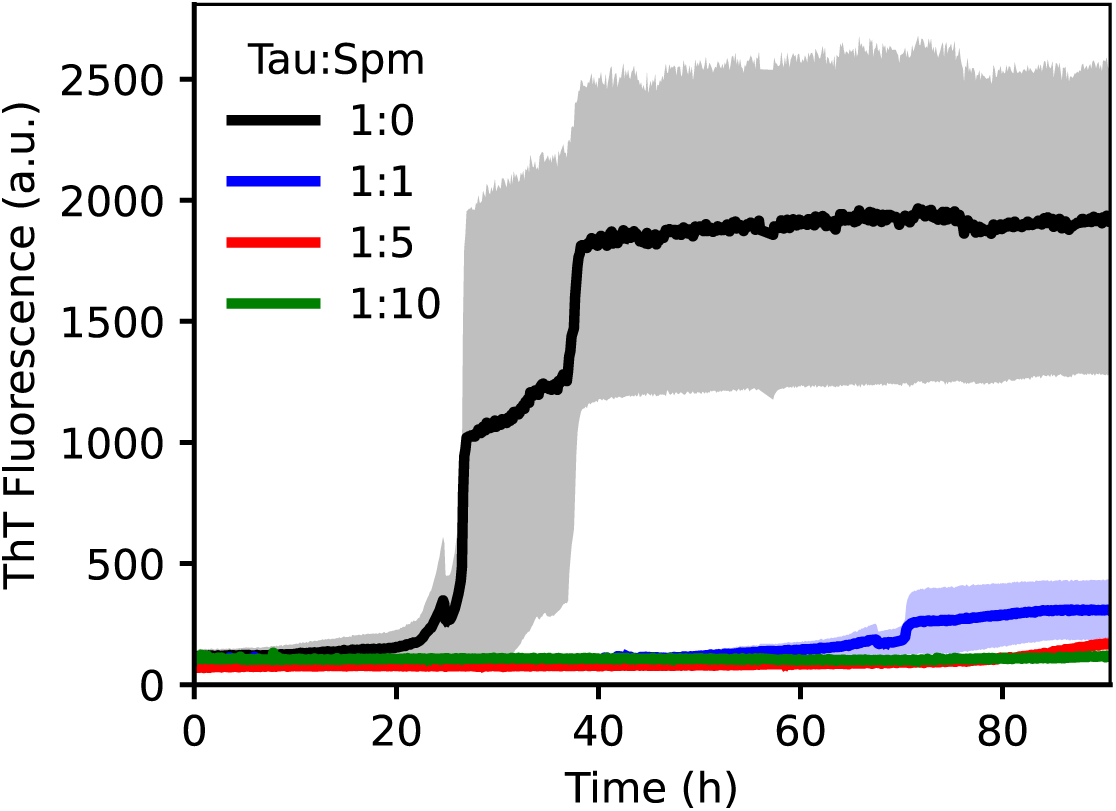
ThT Fluorescence of full-length Tau. For each Tau-Spm ratio, the data shows the average plot along with their standard errors of the mean.

**Figure S9.**
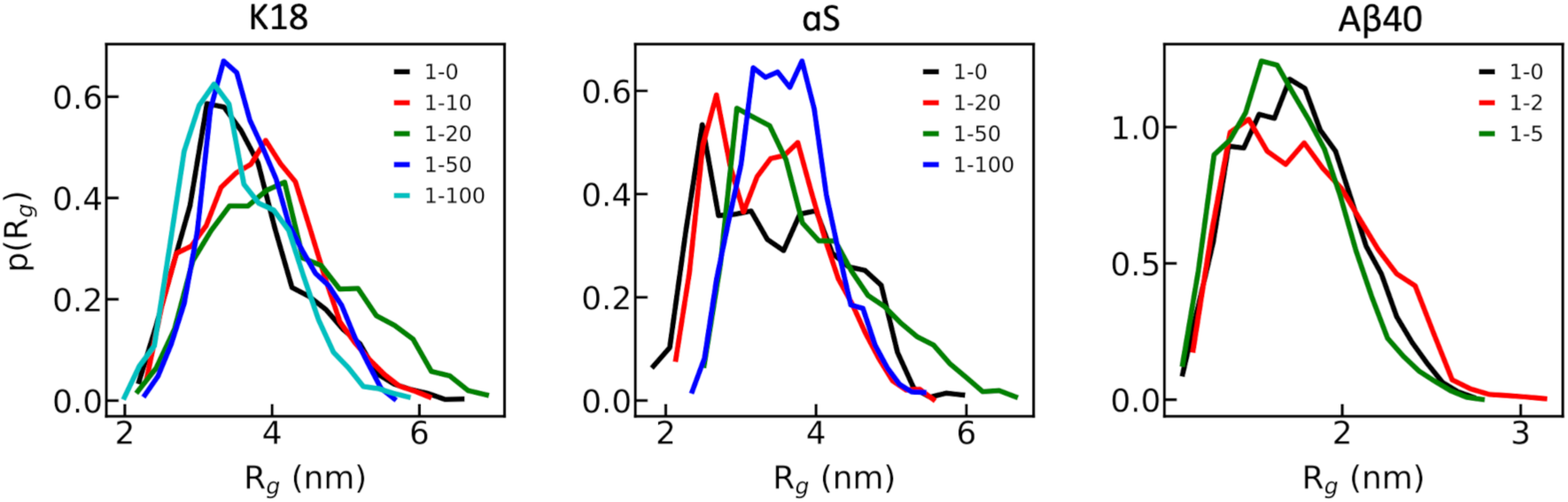
Distribution of *R_g_-s*. The distribution of R_g_s with and without Spm for the three systems.

**Figure S10.**
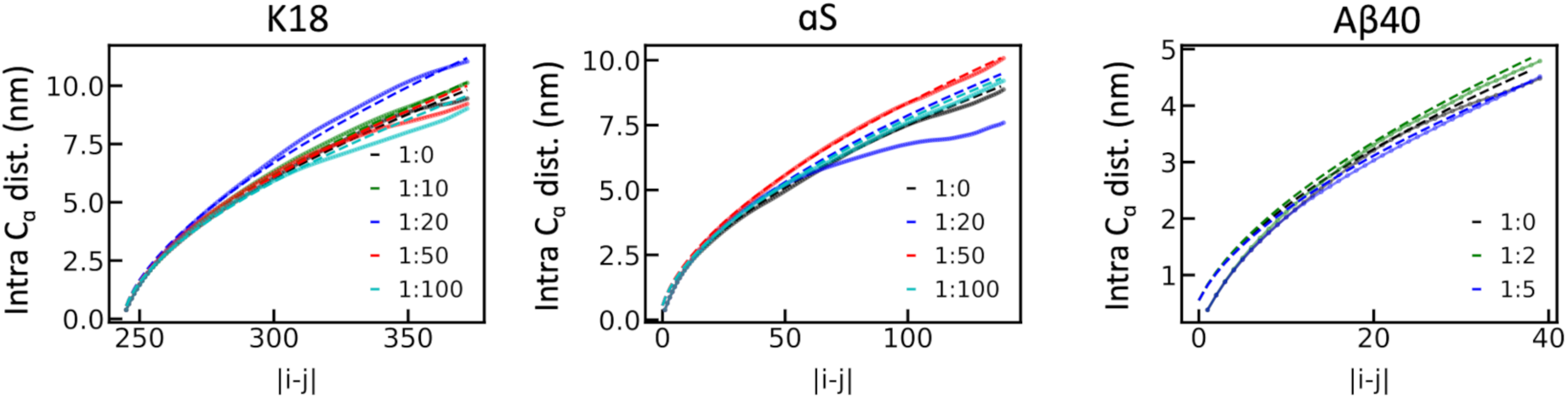
Scaling exponent calculations. The scaling exponents (solid lines) and their fits (dashed lines) for the three systems with and without Spm.

**Figure S11.**
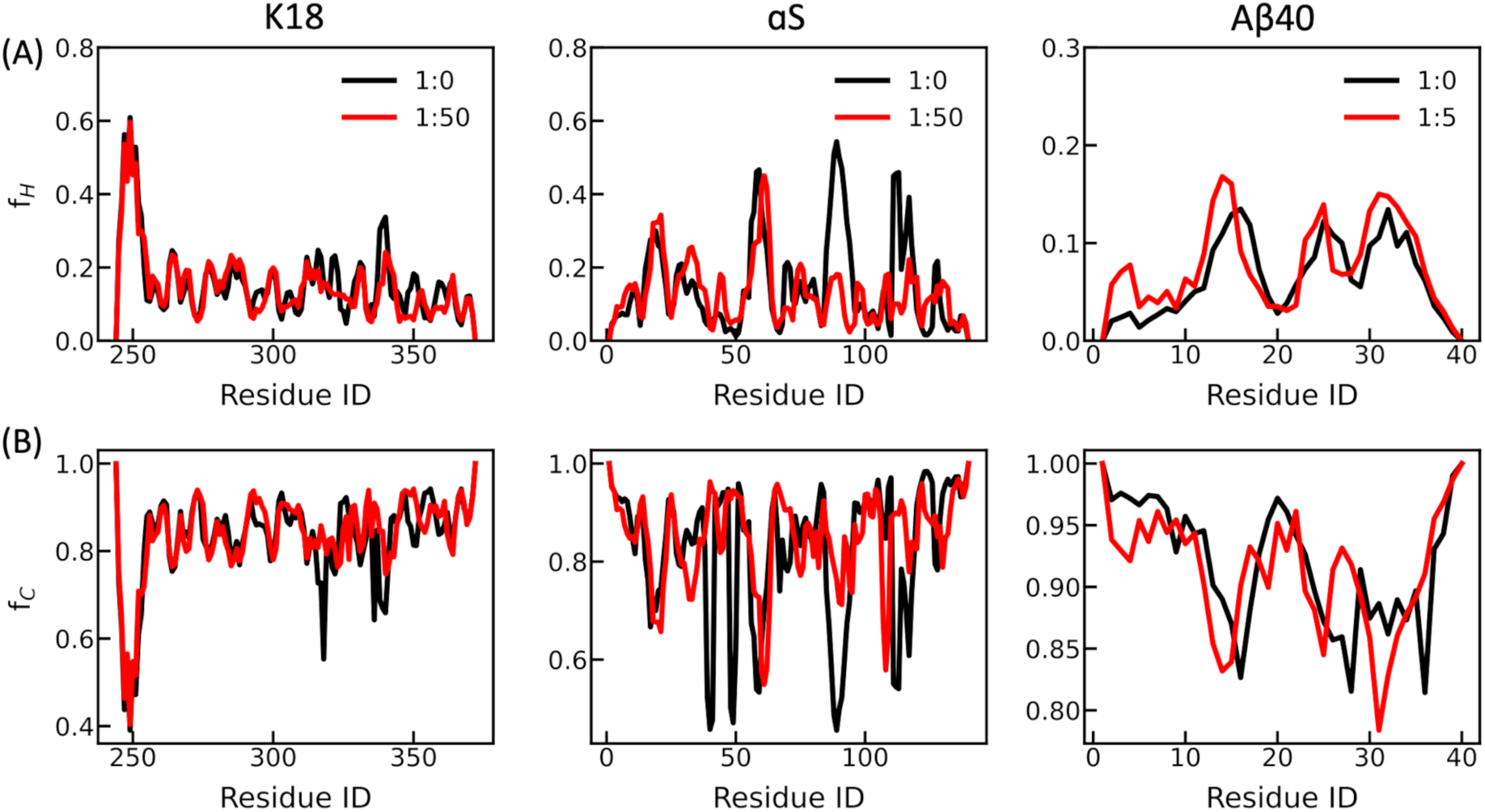
Secondary structure analysis for the three systems. The fraction of helical (H) conformation, *f_H_* (A) and coil (C) conformation, *f_C_* (B) for K18 (left panel), αS (middle panel) and Aβ40 (right panel).

**Figure S12.**
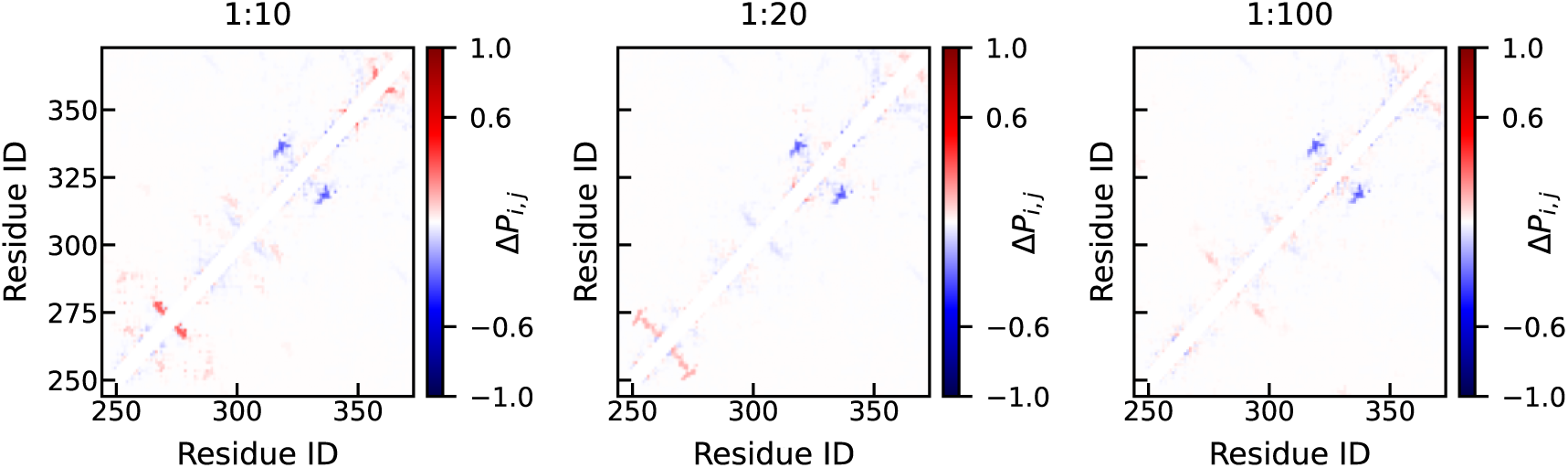
Changes in contact maps for K18 system at various Spm ratios. The changes represent the intra-residue contact probability changes between simulations with and without Spm.

**Figure S13.**
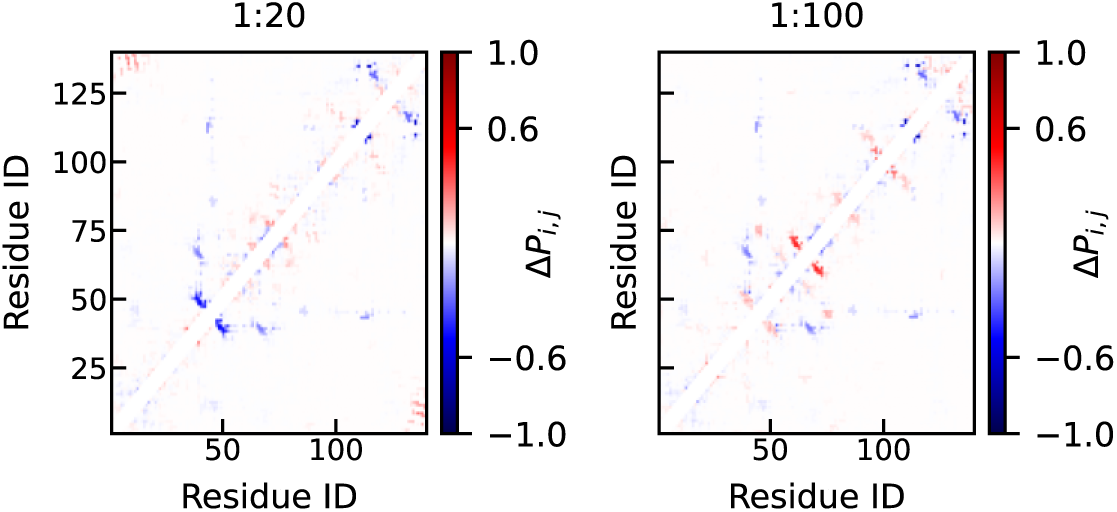
Changes in contact maps for αS system at various Spm ratios. The changes represent the intra-residue contact probability changes between simulations with and without Spm.

**Figure S14.**
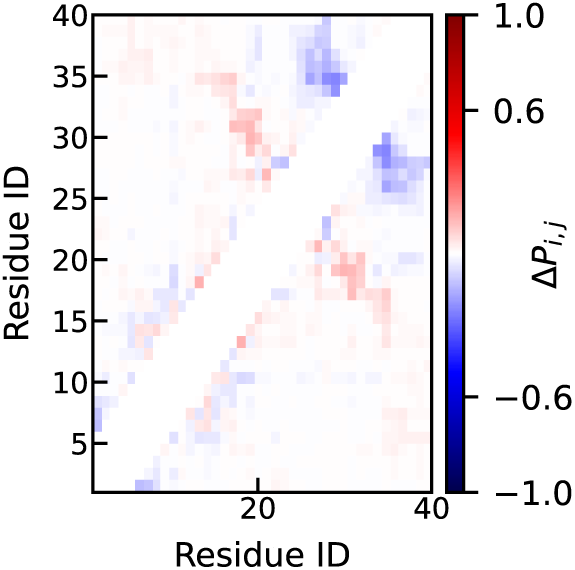
Changes in contact maps for Aβ40 system at 1:2 Spm ratio. The change represent the intra-residue contact probability changes between simulations with and without Spm.

